# A Dynamic NMR Lineshape Simulation Framework for Lipid Diffusion and Membrane Thinning in Bicelles and Nanodiscs

**DOI:** 10.64898/2026.02.24.707804

**Authors:** Sungsool Wi, Ayyalusamy Ramamoorthy

**Affiliations:** National High Magnetic Field Laboratory, Florida State University, 1800 East Paul Dirac Drive, Tallahassee, FL 32310, USA; Department of Chemical and Biomedical Engineering, FAMU-FSU College of Engineering, Florida State University, 2525 Pottsdamer St., Tallahassee, FL 32310, USA; Institute of Molecular Biophysics, Florida State University, 91 Chieftan Way, Tallahassee, FL 32304, USA

## Abstract

Membrane mimetics such as lipid bicelles and nanodiscs have become indispensable platforms for high-resolution structural, dynamical, and functional studies of membrane-associated systems by NMR spectroscopy, cryo-electron microscopy, and X-ray crystallography. In particular, magnetically aligned bicelles and nanodiscs uniquely enable the measurement of anisotropic NMR interactions, providing direct access to membrane geometry, lipid order, thickness, and molecular dynamics. However, the quantitative interpretation of such anisotropic NMR spectra has been hindered by the absence of physically rigorous dynamic models that properly account for the coupled effects of molecular diffusion, orientational distribution, and membrane deformation. Here, we present a comprehensive theoretical framework for the dynamic simulation of ^31^P chemical shift anisotropy and ^14^N quadrupolar NMR lineshapes in bicelles and nanodiscs. The model explicitly incorporates lipid diffusion, orientational distributions on curved membrane geometries, and membrane thinning, enabling physically consistent and quantitatively accurate reproduction of experimentally observed anisotropic lineshapes. Using this framework, we simulate dynamic ^31^P and ^14^N NMR spectra of DMPC/DHPC bicelles and nanodiscs and demonstrate how membrane thinning and lipid diffusion govern the apparent reduction of anisotropic interactions commonly observed upon peptide or protein association. This approach establishes a general physical basis for interpreting anisotropic NMR spectra of aligned membrane mimetics and provides a unified platform for quantitative investigation of membrane structure, dynamics, and membrane-active biomolecular interactions.

## 1. Introduction

The biological cell membrane is a highly dynamic and complex structure that plays essential roles in maintaining cellular integrity, regulating transport, mediating signal transduction, and facilitating interactions between the cell and its environment. In addition to its fundamental physiological functions, the cell membrane is critically involved in the pathology of many diseases, including infectious diseases, neurodegenerative disorders, cancer, and age-related diseases. Achieving a comprehensive understanding of membrane function therefore requires systematic investigation of both the structural organization and dynamic behavior of its constituent components, including lipids, proteins, and other associated molecules. Significant progress has been made in elucidating atomic-resolution structures of membrane proteins and characterizing the biophysical properties of lipid bilayers using a variety of experimental and computational approaches. [1] [2] [3] [4] [5] Despite these advances, the amphipathic nature of membrane components and the compositional and functional complexity of biological membranes continue to pose substantial challenges for *in vitro* studies. Native cell membranes are heterogeneous and highly dynamic, making them difficult to reproduce accurately under controlled experimental conditions. To address these challenges, several different types of membrane mimetic systems have been developed to approximate key features of biological membranes while enabling the application of biophysical and atomic-resolution structural techniques. Commonly used membrane mimetics include lipid vesicles, bicelles, nanodiscs, membrane blebs, and exosomes, each of which offers a unique balance between experimental tractability and physiological relevance. [6] [7] [8] [9] [10] In contrast, detergent micelles, although used in membrane protein purification and structural studies, are generally regarded as suboptimal membrane mimetics. Each membrane mimetic system presents distinct advantages and limitations with respect to size, lipid composition, curvature, stability, and compatibility with analytical methods. Among these systems, bicelles and nanodiscs have emerged as particularly powerful tools for high-resolution structural and dynamical studies of membrane-associated proteins. [11] [12] [13] [14] [15] [16] Their ability to provide a relatively native-like lipid bilayer environment, combined with their tunable size and composition, makes them better suited for probing protein–lipid interactions by NMR spectroscopy. Consequently, bicelles and nanodiscs are widely employed to investigate the structure, dynamics, and mechanisms of action of membrane-active peptides, including amyloid peptides, antimicrobial peptides, toxins, and viral fusion peptides. These studies have provided critical insights into how such peptides interact with lipid bilayers, disrupt membrane integrity, or induce membrane remodeling, thereby advancing our understanding of both normal membrane-associated processes and membrane-related disease mechanisms.

Bicelles and nanodiscs are discoidal nanoparticles composed of a planar lipid bilayer patch encased by a stabilizing belt. [11] [13] This belt can be formed from detergents, short-chain lipids, membrane scaffold proteins (MSPs), amphipathic peptides, synthetic polymers, or saponin molecules. [10] [15] The belt stabilizes the exposed hydrophobic edges of the lipid bilayer, allowing these assemblies to remain soluble and structurally well defined in aqueous environments. The geometry and physical properties of bicelles and nanodiscs are highly sensitive to experimental conditions, including temperature, lipid composition, the molar ratio of lipids to belt-forming components, and the presence of divalent metal ions. Variations in these parameters can significantly influence particle size, bilayer thickness, and overall morphology. For example, at elevated temperatures, discoidal nanoparticles may collide and fuse, leading to the formation of larger bilayer slabs or extended lamellar structures. On the other hand, under extreme conditions, such as very high temperatures or unfavorable lipid-to-belt ratios, these assemblies can fragment to form smaller vesicles, micelles, or mixed aggregates of lipids and belt components. This tunable structural behavior makes bicelles and nanodiscs versatile membrane mimetics for biophysical and structural studies, but it also necessitates careful control and characterization of experimental conditions to ensure reproducibility and physiological relevance.

Small-sized bicelles and nanodiscs, that undergo rapid isotropic tumbling on the NMR timescale, are widely used in solution NMR studies to investigate the structure, dynamics, and interactions of embedded peptides and proteins. [17] [18] [19] [20] Their fast rotational diffusion leads to narrow resonance linewidths, enabling high-resolution spectral analysis and detailed characterization of membrane-associated biomolecules. In contrast, large-sized bicelles/nanodiscs that are capable of aligning in the presence of an external magnetic field are commonly employed in solid-state NMR experiments. [21] [22] [23] These aligned bicelle/nanodisc systems enable the investigation of the structure and dynamics of all membrane components, including lipids, proteins or peptides, cholesterol, and other associated molecules. [24] [25] They also provide a powerful platform for probing the effects of various factors on lipid bilayer properties, such as membrane phase behavior, molecular ordering and disordering, bilayer thickness, and overall membrane stability. [26]

Solid-state NMR (ssNMR) experiments under static conditions (and in some cases under magic-angle spinning (MAS)) commonly utilize nuclei such as ^31^P, ^14^N, ^2^H (for ^2^H-labeled molecules), ^13^C, and ^1^H. In particular, ^31^P, ^14^N, and ^2^H spectral lineshapes obtained from magnetically-aligned bicelles or nanodiscs are routinely compared with powder-pattern lineshapes from unoriented lipid vesicles to determine the macroscopic orientation of individual molecular components within the membrane. [27] For example, unaligned lamellar-phase DMPC large unilamellar vesicles (LUVs) exhibit a characteristic ^31^P chemical shift anisotropy (CSA) powder pattern spanning approximately −14 to +30 ppm (referenced to H□PO□ at 0 ppm). In contrast, DMPC lipids incorporated into bicelles or nanodiscs display a narrow isotropic 31P resonance at low temperatures, below the gel-to–liquid-crystalline phase transition temperature (Tm). Upon increasing the temperature above Tm, a narrow anisotropic resonance appears at the perpendicular edge of the CSA (approximately −14 ppm), indicating uniform macroscopic magnetic alignment of the lipid bilayers with their normals oriented perpendicular to the magnetic field axis. Notably, this orientation can be flipped by 90° resulting in bilayer normals aligned parallel to the magnetic field through the introduction of paramagnetic dopants, providing additional experimental flexibility for structural analysis. In contrast, ^14^N (spin = 1) nuclei present in the choline moiety of the phosphocholine headgroup exhibit a quadrupolar coupling powder pattern in DMPC large unilamellar vesicles (LUVs) that is substantially reduced due to motional averaging of the lipid molecules, as well as the near-tetrahedral symmetry of the nitrogen environment. In magnetically aligned bicelles or nanodiscs, however, the ^14^N spectra display narrow doublet resonances corresponding to the perpendicular edge of the quadrupolar powder pattern. Upon a 90° flip of the macroscopic alignment by paramagnetic doping, these doublet peaks shift to the parallel edge of the powder pattern, reflecting the change in bilayer orientation relative to the external magnetic field. Similarly, ^2^H-labeled lipids are commonly used to measure ^2^H quadrupolar couplings, providing quantitative information on lipid structure, segmental order, and molecular mobility within lipid bilayers. The ability to exploit motionally averaged anisotropic interactions, such as ^31^P CSA and ^14^N and ^2^H quadrupolar couplings, to determine the orientation and dynamics of individual components of lipid membranes is a powerful approach with broad applicability. These measurements enable detailed evaluation of factors that influence membrane integrity, stability, thickness, and the degree of lipid order or disorder.

Owing to these capabilities, ^31^P and ^14^N solid-state NMR are extensively used to elucidate the mechanisms of action of membrane-active agents, including antimicrobial peptides, amyloid peptides, fusogenic peptides, and toxins. However, accurate interpretation of ^31^P and ^14^N NMR parameters critically depends on the lipid bilayer model employed to simulate the experimentally observed spectra. Structural features such as the presence of lipids in the rim region and toroidal pores of bicelles, lipid domains present in multicomponent bicelles or nanodiscs, the influence of belt-forming molecules in nanodiscs, and perturbations introduced by additional components (e.g., peptides, proteins, cholesterol, or small-molecule drugs) can substantially complicate spectral interpretation. As the use of bicelles and nanodiscs continues to expand toward increasingly complex and biologically relevant membrane systems, there is growing interest in the development of comprehensive and physically realistic models capable of accurately reproducing experimental NMR observables. In this study, we define the geometry of bicelles and nanodiscs and develop a mathematical framework that accounts for the distinct orientations and diffusive behaviors of lipid populations within these assemblies. Using this framework, we accurately simulate ^31^P and ^14^N NMR spectra of lipids under a range of experimental conditions. Simulated spectra demonstrating the validity and robustness of the model for several bicelle and nanodisc systems are presented and, wherever possible, directly compared with corresponding experimental results.

## 2. Simulation Methodology

Anisotropic solid-state NMR (ssNMR) spectra of ^31^P and ^14^N (or ^2^H) nuclei are widely used to characterize anisotropically disordered, oriented, and aligned lipid assemblies. [28] [29] [30] [31] The anisotropic ssNMR lineshapes arise from the ^31^P CSA and ^14^N (or ^2^H) QC interactions defined in the laboratory frame, as [32]

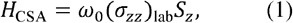

and

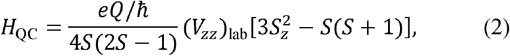

respectively. Here, *ω*_0_ = *γB*_0_ is the Larmor frequency, *S*_*z*_ is the angular momentum operator component along the applied magnetic field *B*_0_, and ***σ***_lab_ and ***V***_lab_ are the 3 ×3 CSA and electric-field-gradient (EFG) tensors in the laboratory frame. The quadrupolar interaction is commonly expressed as *eQV*/ *ħ* representing the electrostatic interaction between the nuclear quadrupole moment *eQ*/ *ħ* and the EFG tens *V*. For lipid molecules assembled into supramolecular membrane structures, the CSA and QC interaction tensors provide direct information on the angular distributions of the corresponding nuclear sites relative to the static magnetic field *B*_0_. Consequently, the resulting ssNMR lineshapes directly reflect the orientational and geometrical distributions of lipid molecules with respect to *B*_0_. In the molecular-specific principal axis system (PAS), only the diagonal tensor elements are non-zero. For the CSA interaction, the standard tensor conventions are [32] [33][34]

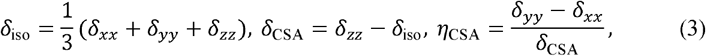

while for the quadrupolar interaction, [35] [32] [33] [36]

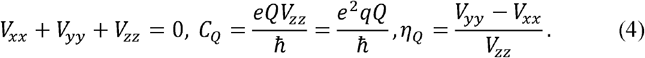

Here, *δ*_CSA_ and *C*_*Q*_ denote the magnitudes of the CSA and quadrupolar tensors, respectively, and *η*_*λ*_(*λ* =CSA, QC) is the asymmetry parameter. The chemical shift is expressed using *δ* rather than *σ*, assuming referencing to a standard compound via *δ* =*σ*_ref_ −*σ*.

Under sufficiently hydrated conditions, a two-tailed phospholipid molecule in a bilayer undergoes rapid uniaxial rotation on a nanosecond timescale about its molecular long axis, which is collinear with the local bilayer normal. [37] This fast uniaxial motion leads to motional averaging of the intrinsic second-rank interaction tensors, resulting in axially symmetric, motionally averaged CSA and QC tensors for the ^31^P and ^14^N (or ^2^H) sites, respectively. For example, rapid uniaxial rotation of a lipid molecule about its acylchain axis—assumed here to be collinear with the *δ*_*yy*_ principal axis of the intrinsic ^31^P CSA tensor—yields motionally averaged CSA tensor elements

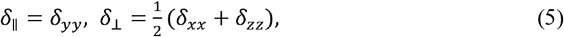

with *η*_CSA_ = 0. [37] As a result, lipid assemblies such as liposomes, bicelles, toroidal pores, and related membrane morphologies exhibit a significantly narrowed apparent ^31^P CSA powder pattern compared to that of the intrinsic CSA tensor. An analogous motional averaging mechanism applies to the quadrupolar couplings of ^14^N nuclei in the hydrophilic choline headgroup and of ^2^H nuclei incorporated into the hydrophobic acyl chains of perdeuterated lipids, yielding axially symmetric, motionally averaged QC tensors. The ratio of the magnitudes of the motionally averaged CSA or QC tensors to their intrinsic tensor magnitudes defines an order parameter, which quantitatively characterizes the degree of lipid motional mobility within the bilayer.

The observed anisotropic resonance frequency of a nuclear site in ^31^P, ^14^N, or ^2^H ssNMR experiments depends explicitly on the orientation of the corresponding interaction tensor relative to *B*_0_, while employing the motionally averaged, apparent uniaxial tensor elements. For lipid molecules distributed within bicelle assemblies, we define the bilayer normal vector of the flat discoidal region as the reference alignment axis. Although not addressed in this manuscript, in mechanically oriented lipid samples prepared between glass plates, *n* is assumed to be collinear with the plate normal. The anisotropic resonance frequencies in the rotating frame (equivalently, the laboratory frame) are given by

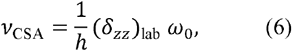

for ^31^P CSA, and

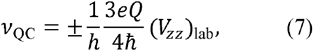

for ^14^N (or ^2^H) QC, where the apparent, motionally averaged interaction tensors are employed. These expressions correspond to the 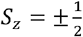 transition for ^31^P and the *S*_*z*_ = ±1 ↔ 0 transitions for ^14^N (or ^2^H). The spatial tensors *δ*_lab_ and *eQV*_lab_/ *ħ* are related to their counterparts in the motionally averaged PAS via coordinate transformations. When the bilayer normal is parallel to the magnetic field (n ∥ *B*_0_) the transformation from the PAS to the laboratory frame is described by the Euler angle set Ω ={*ξ*=0°, *θ, ϕ*= 0°} reflecting axial symmetry (*ξ*=0°) and the irrelevance of rotation about *B*_0_ (*ϕ* =0°).This transformation is expressed as

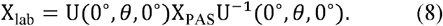

The PAS tensors are given by

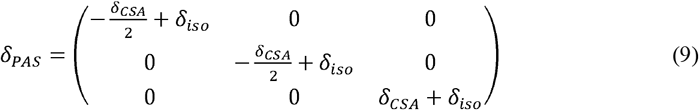

and

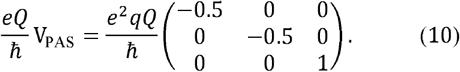

The rotation matrix U(*α, β, γ*) defined by the Euler angles Ω ={*α, β, γ*}, is

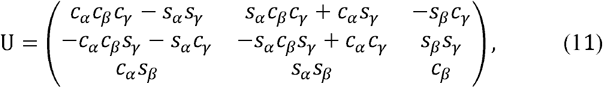

where *c* ≡cos and *s* ≡ sin. [32] Large bicelles are known to spontaneously align in magnetic fields with their bilayer normal perpendicular to *B*_0_ (n ⊥ *B*_0_). For this native alignment, an additional coordinate transformation from the bilayer-normal frame to the laboratory frame is required, using the Euler angle set {*ϕ*,90°,0°}: [38] [39]

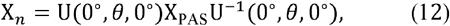

The n ⊥ *B*_0_ orientation corresponds to the native magnetic alignment of bicelles, whereas the n ∥ *B*_0_, configuration can be achieved by rotating the bicelles by 90°, typically through the addition of paramagnetic alignment agents such as lanthanide chelates.

The n ⊥ *B*_0_orientation corresponds to the native magnetic alignment of bicelles, whereas the n ∥ *B*_0_, configuration can be achieved by rotating the bicelles by 90°, typically through the addition of paramagnetic alignment agents such as lanthanide chelates.

For lipid molecules distributed over curved membrane surfaces, the anisotropic resonance frequency *v*_*λ*_ depends explicitly on the local membrane geometry. The resulting ssNMR lineshape is obtained by integrating over the Euler angle set Ω = (0°,*θ, ϕ*) with 0 ≤ *θ* ≤ *π* and 0 ≤ *θ* ≤ *π* and 0 ≤ *ϕ* ≤ 2*π*:

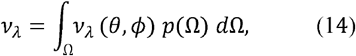

where *p*(Ω) is the probability density function proportional to the infinitesimal surface area element and is referred to as the anisotropic NMR lineshape factor. For randomly oriented lipids or spherical liposomes, *p*(Ω) = sin *θ*, whereas for planar or cylindrical geometries, *p*(Ω)=1. The specific forms of *p*(Ω) for bicelles (nanodiscs), toroidal pores, and membrane-thinning geometries are described in the Results section. All ssNMR lineshape simulations were performed by numerically propagating the total magnetization while sampling powder orientations over *θ* and *ϕ*. A Lorentzian line-broadening of 50 Hz was included in all simulations. Lipid lateral diffusion was explicitly incorporated to account for dynamic averaging effects. For bicelle simulations, realistic molecular counting was achieved by specifying the bicelle size and geometry based on the bilayer thickness and the *q*-factor, defined as *q* = mol(DHPC)/mol(DMPC). Following time-domain signal generation, Fourier transformation of the simulated free induction decay yielded the final motionally averaged anisotropic ssNMR lineshapes.

All simulations were carried out using a home-built MATLAB program. The codes used to simulate ^31^P and ^1^□N (or ^2^H) ssNMR lineshapes for bicelle and membrane-thinning models are provided in the Supporting Information and are made freely available to the community. Owing to its generality, this framework is readily extendable to other anisotropically ordered membrane assemblies and provides a powerful quantitative tool for linking ssNMR lineshapes to membrane geometry, molecular population statistics, and lipid dynamics in complex soft-matter systems.

## 3. Results and Discussion

Small-angle neutron scattering (SANS), [40] small-angle X-ray scattering (SAXS), [41] and combined NMR/SAXS [42] studies have established that DMPC/DHPC bicelles adopt oblate ellipsoidal shapes rather than idealized circular disks. In this context, anisotropic solid-state NMR spectra of ^1^□N (or ^2^H) and ^31^P nuclei are uniquely powerful probes of lipid supramolecular organization, as they report directly on the molecular-level orientational distributions of lipid headgroups and acyl chains. [43] [44] [45] Consequently, these nuclei have been used extensively to characterize bicelle structures and their interactions with membrane-active peptides and proteins. In this study, we employ simplified geometric models to describe the distribution of lipid and belt-forming molecules in bicelles and nanodiscs, and torus as illustrated in Figure 1. Although the model shown in (A) is applicable to both bicelles and nanodiscs systems, the belt-forming molecules in nanodiscs differ substantially from those in bicelle and do not contribute to the commonly observed ^31^P and ^14^N NMR spectra. Therefore, to develop a mathematical framework for interpreting experimentally observed ^31^P and ^14^N solid-state NMR spectra, we focus on bicelles, while noting that the approach presented here is equally applicable to lipid molecules present in a nanodisc. In a typical DMPC:DHPC bicelle, DMPC lipids form a bilayer patch that is surrounded by DHPC detergent molecules. Depending on sample conditions, DHPC molecules may also form toroidal pores (Fig. 1B) within the planar bilayer region. Consequently, DMPC molecules are expected to exhibit a uniform uniaxial orientation within the bilayer, whereas DHPC molecules in the rim and toroidal pores are expected to adopt a toroidal distribution. The presence of membrane-interacting molecules such as proteins, peptides, cholesterol, or drugs can perturb the orientations, axial rotation and diffusion of DMPC and DHPC molecules. To account for these effects, we develop a general theoretical framework capable of accurately simulating ^31^P and ^14^N solid-state NMR spectra, enabling determination of molecular orientations and diffusion dynamics in bicelles. Simulated spectra under various conditions are presented and, where applicable, compared with experimental results.

**Figure 1.**
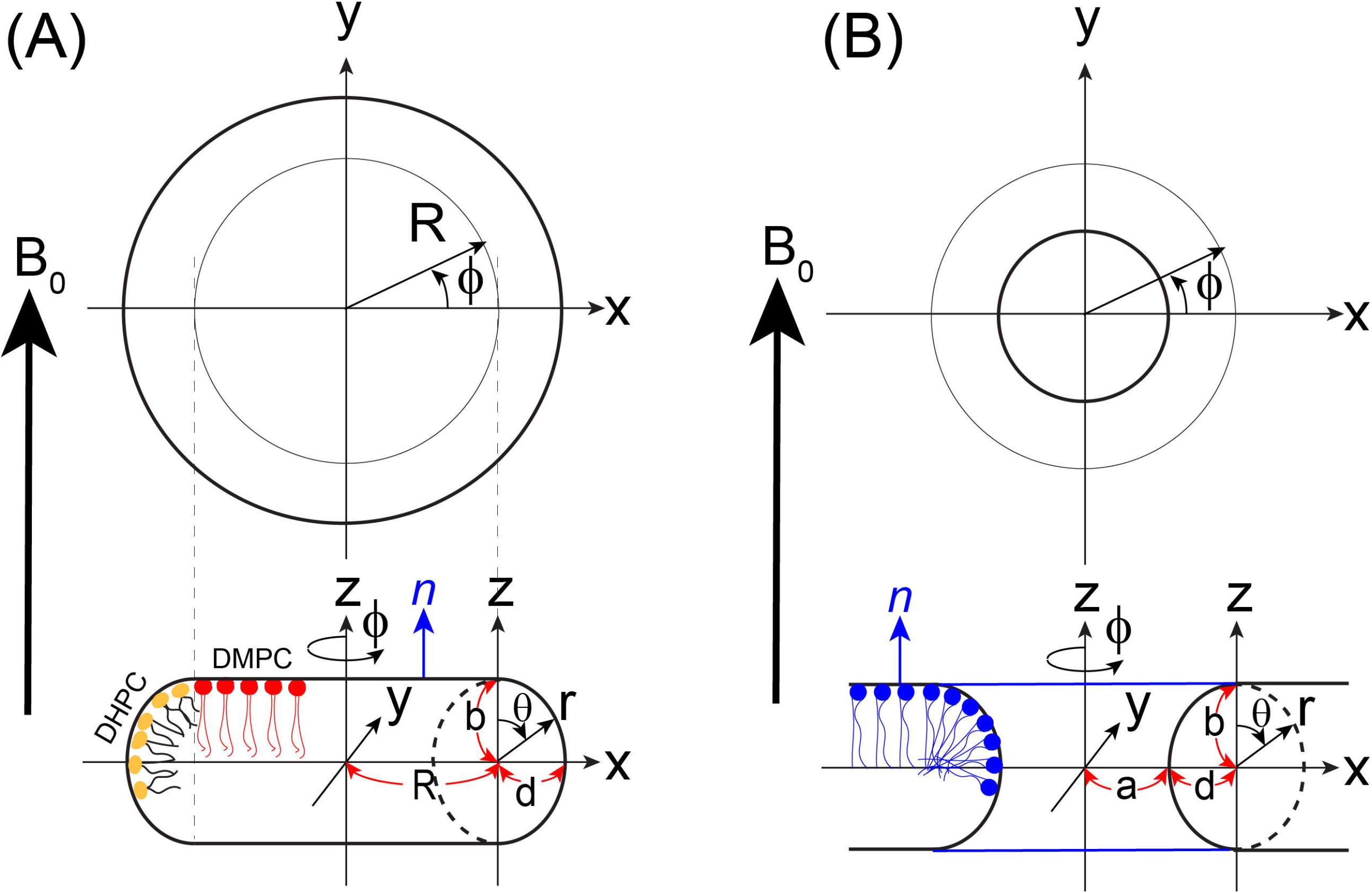
Simplified geometric models used for solid-state NMR lineshape analysis of (A) bicelles and (B) toroidal pores. Lipids are assumed to populate both the flat bilayer regions and the curved rim or pore surfaces. The geometries shown correspond to the case in which the bilayer normal n in the flat region is parallel to the external magnetic field (n ∥B□). The perpendicular orientation (n⍰ B□) can be treated by applying an additional tensor transformation to the n ∥ B□ configuration, as described in the text. For the bicelle model (A), the radius *R* of the flat bilayer region is determined by the *q*-value, defined as the molar ratio *q* = *m*(DMPC)/*m*(DHPC). For the toroidal pore model (B), the central region of radius *a* is devoid of lipids, where *a* corresponds to the narrowest pore diameter. In both models, the curved regions (bicelle rim and pore surface) are approximated by an elliptic cross-section, enabling unified treatment of lipid orientation and curvature effects in the NMR simulations.

### 3.1. Bicelle and Torus Geometries for NMR Lineshape Simulations

Lipid molecules distributed on the bicelle and toroidal pore geometries give rise to characteristic ^31^P, ^2^H, and ^14^N NMR lineshapes. To accurately reproduce these features, we analytically develop geometrical models for describing both bicelle and torus pore, and further incorporate a description of lateral molecular diffusion on these surfaces based on Fick’s second law to take into account the dynamic nature of molecules distributed on these curved surfaces. Fig. 1A depicts the case where the lipid bilayer normal (n) is oriented parallel to the external magnetic field (n//B_0_). Using this flat discoid geometry, the two planar faces of the discoid are employed to represent the lipid bilayer regions of a bicelle, emulating lipid distributions within the discoidal planes. The continuously curved outer surface enclosing the flat discoid—generated by a full 360° rotation about the *z*-axis passing through the center of the discoid and perpendicular to the *x–y* plane—is treated as the rim region of the bicelle, where molecules experience increased curvature relative to the central bilayer region. This construction fully specifies the bicelle geometry used in the simulations. Independently, a similar type of the geometric framework is used to model a toroidal pore (Fig. 1B). [38] [39] Again, upon a full 360° rotation of an elliptic circle defined in the xz-cross section about the *z*-axis, this profile generates the inner curved surface of a toroidal pore. It should be emphasized that, while the bicelle geometry is completely defined by this construction, the toroidal-pore model uniquely specifies only the inner curved surface. Because the geometry is open outwardly, the flat bilayer regions associated with the pore cannot be determined within this simple geometric framework and therefore remain undefined. More detailed descriptions of these models, including the distances and angular parameters used in the construction are provided below.

#### 3.1.1. Bicelle geometry and surface parameterization

In the bicelle simulation, molecules are distributed over both the curved rim region and the two flat discoid faces representing the bilayer surfaces. For lipids located on the flat regions, the bilayer normal *n* is explicitly included to account for their orientational dependence. The bicelle geometry defined in Fig. 1A is described by the following parametric equations:

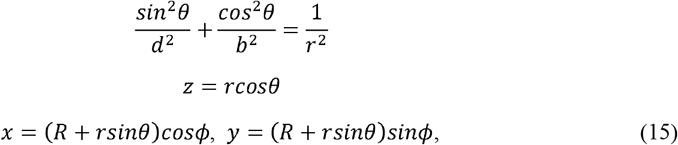

where an elliptical arc is used to represent the rim region at the xz-cross section. This elliptical cross section provides flexibility in modeling the rim structure and is characterized by a major axis *b* and a minor axis *d*, whose roles may be interchanged depending on the specific geometry considered. In the present model, *b* corresponds to half of the bilayer thickness (2*b*), which is typically ∼40 Å for phospholipid bilayers composed of lipids such as POPC or DMPC. The parameter *R* can be specified according to the q-ratio (*m*(DMPC)/*m*(DHPC)) as will be explained later. Lipids located in the flat disc region give rise to a single resonance line, the position of which is determined by the orientation of the bilayer normal n relative to the external magnetic field B□ (i.e., n ∥ B□, n ⍰ B□, or an arbitrary tilt angle).

To compute the NMR lineshape contributions from lipids distributed along the curved rim, the relevant surface element is evaluated. Since *r*(*θ*) is a dependent variable, the Jacobian determinant *J*= *det* (*∂* (*x, y, z*) /*∂*(*r, θ, ϕ*)) is replaced by the surface element [38] [39]

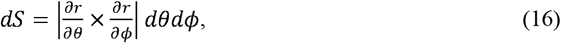

which yields

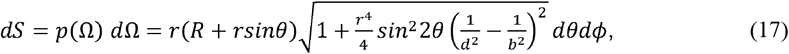

With

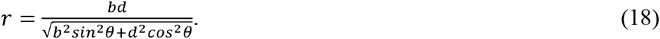

#### 3.1.2. Toroidal pore geometry

An analogous geometric framework (Fig. 1B) is used to simulate lipids distributed along the inner surface of a toroidal pore. In this case, the surface outlined by the inner curved surface must be considered [46] [39] [38]

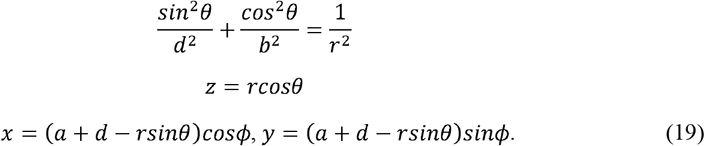

The corresponding surface element required for NMR lineshape simulations is

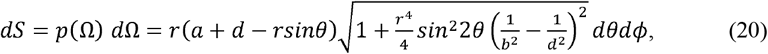

with

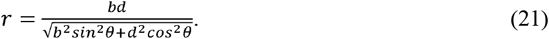

#### 3.1.3. Angular dependence and lipid orientation

In both geometries, the NMR lineshape factor associated with the azimuthal angle □ is unity. The θ-dependent contribution to the surface weighting is given by

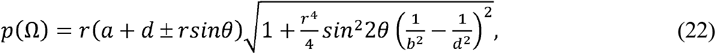

where the plus sign applies to the bicelle rim (a+d = R) and the minus sign applies to the toroidal pore. In the special circular case (*b = d*), this expression simplifies to *b(R + b sin θ)* for the bicelle rim and *b(a + b − b sin θ)* for the toroidal pore. [46] In this limit, the lipid normal along the curved surface is always orthogonal to the surface tangent. However, when *b* ≠ *d*, this orthogonality condition is no longer automatically satisfied. To enforce proper alignment of lipid normals with the curved surface, a modified angular variable θ′ is introduced: [38]

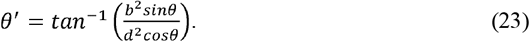

Accordingly, θ′ is used in place of θ in the geometric description when determining lipid orientations on curved surfaces. In the present manuscript, NMR lineshape simulations are performed for bicelle lipid distributions. The toroidal pore case, while formulated here for completeness, has been reported previously. [39] [38]

### 3.2. Consideration of Lateral Diffusive Motion of Lipids

For the simulation of ^31^P, ^2^H, or ^1^□N NMR spectra of lipid supramolecular assemblies, including lipid bilayer, bicelles, toroidal pores, and dimples, geometry alone is generally insufficient to fully describe the observed spectra. Two additional dynamic effects must be taken into account in the simulations of lipid assemblies. First, under sufficiently hydrated conditions, individual lipid molecules within a bilayer undergo rapid uniaxial rotational motion about the bilayer normal. As a result of this fast motion, the ^31^P chemical shift anisotropy (CSA) and 14N or 2H quadrupolar interactions are motionally averaged to result in axially symmetric tensors with an asymmetry parameter *η* =0. In practice, this effect is directly incorporated into simulations by using pre-averaged, axially symmetric CSA or quadrupolar tensor parameters. The order parameter of the lipid site is obtained from the magnitudes of these motionally averaged tensor parameters. Second, lipid molecules also undergo rapid lateral diffusion over the surface of the supramolecular lipid assembly. Unlike the uniaxial rotational averaging, this lateral diffusive motion cannot be captured simply by using averaged tensor parameters. Instead, it must be treated explicitly through an appropriate dynamic diffusion model.

To account for the lateral diffusive motion of lipids on the surface of a lipid supramolecular assembly, the time evolution of the magnetization must satisfy Fick’s diffusion law (or the second law), [30] [47] [38]

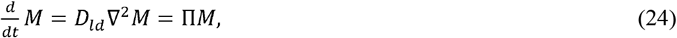

where *D*_1d_ is the lateral diffusion coefficient, *M* is the magnetization, and Π denotes the diffusion rate describing how the magnetization changes at a specific point over time. The Laplacian in a general curvilinear coordinate system *(q*_*1*_, *q*_*2*_, *q*_*3*_) is given by

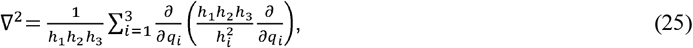

where *h*_*i*_ (*q*_*i*_) are the corresponding scale factors. Then Eq. (24) can be discretized and approximated as

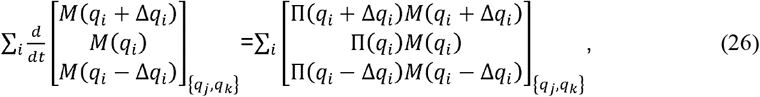

where the diffusion rate evaluated at the displaced coordinates is defined as

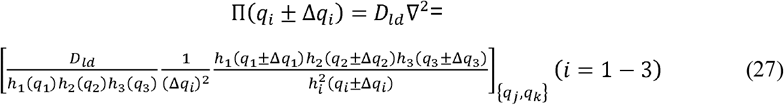

and the rate at the central grid point is approximated by

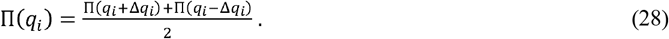

Here, Δ*q*_*i*_=*q*_*i*+1_ − *q*_*i*_ denotes the finite step size along the *q*_*i*_ coordinate, and the subscripts *q*_*j*_,*q*_*k*_ indicate that the remaining coordinates are held fixed during the evaluation.

Among the three coordinates (r, θ, □) defining the bicelle discoid, lipid motions along the r-direction are excluded because it is orthogonal to the surface on which lateral diffusion takes place. Thus,

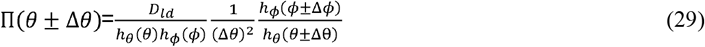

and

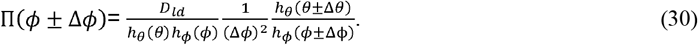

Explicit expressions for the Π (*θ* ±Δ *θ*) and Π(*ϕ* ± Δ *ϕ*) for the bicelle model are provided in the Supporting information. Note that the product of the surface distance scale factors, *h*_*θ*_ (*θ*) *h*_*ϕ*_ (*ϕ*) corresponds to the NMR lineshape weighting factor Ω defined previously.

If the position of a ^31^P, ^14^N, or ^2^H site in a lipid molecule is described by an angular coordinate set (*θ, ϕ*), its corresponding anisotropic NMR frequency is de termi ned by this angle set with respect to the static magnetic field *B*_0_ and can be written as *v*(*θ, ϕ*). To incorporate lateral diffusion,the orientational (*θ, ϕ*) space is discretized into a finite set of grid points (*θ*_*i*,_*ϕ*_*i*_) = (*n*_*i*_ Δ*θ,m*_*i*_Δ *ϕ*), where *ni* and.*mi* are integers. Each discrete orientation is thus associated with a specific anisotropic frequency *v*_*i*_ (*θ*_*i*_, *ϕ*_*i*_). We assume that a lipid occupying agiven orientational state (*θ*_*i*_, *ϕ*_*i*_) can migrate to its nearest neighboring states through lateral diffusion on the supramolecular surface. This process is described by diffusive translation between adjacent grid points along the *θ* and *ϕ* directions,

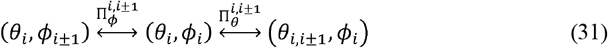

where 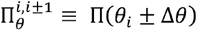 and 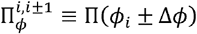 denote the diffusion-induced transition rates between neighboring orientation sites in the lipid distribution along the *θ* and *ϕ*,coordinates, respectively. In this manner, lateral lipid diffusion on the curved supramolecular surface is modeled as a sequence of successive random walks in orientational space, leading to time-dependent redistribution of the anisotropic frequencies *v*_*i*_ (*θ*_*i*_, *ϕ*_*i*_) that contribute to the observed NMR lineshape.

For the simpler case in which n ∥ B□, rotation about the azimuthal angle *ϕ* around the *z*-axis (∥ B□) does not affect the NMR lineshape. Therefore, only the θ-dependence (θ is decomposed into nΔθ steps) is considered in the NMR exchange simulations arising from lateral lipid diffusion. Assuming stepwise lateral diffusive jumps between nearest neighbors, the corresponding tridiagonal exchange matrix is given by

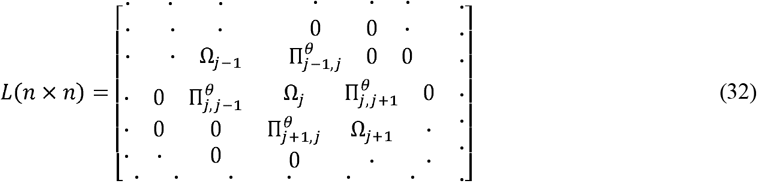

where Ω*j* (*j* =1,2, …,*n*) are diagonal elements given by

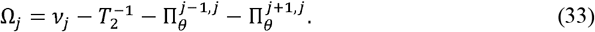

Here, *v*_*j*_ and 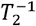 are the orientation-dependent anisotropic NMR frequency and the spin-spin relaxation rate at site *j*, respectively, and it is assumed that 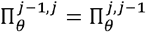 and 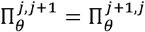. The time evolution of the transverse magnetization M^+^ arising from this exchange model can be calculated using the Bloch–McConnell differential equation [48] [49]

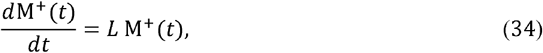

where M+ (*t*) is an (*n* × 1) column vector representing the transverse magnetization intensities at all lattice points. In matrix form, the solution of Eq. (34) is given by

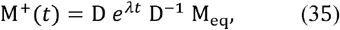

where M_eq_ denotes the equilibrium population of the lattice points, which is weighted by the NMR lineshape factors for the lipid geometry under consideration, and D and *λ* are the eigenvectors and eigenvalues of the exchange matrix *L*, respectively. Summation of all components of M+(*t*), followed by Fourier transformation, yields the motionally averaged exchange NMR spectrum arising from lateral lipid diffusion on the surface of a curved membrane bilayer for n ∥ B□ case.

For the more complex case of n ⍰ B□, a coordinate transformation is required, as specified in Eqs. (12-13), in which both the polar angle θ and the azimuthal angle □ are treated simultaneously. The θ and □ dependencies in the simulation are discretized into *n* steps of Δθ and *m* steps of Δ□, respectively, where Δθ and Δ□ denote the angular step sizes. This discretization leads to an *(nm) × (nm)* supermatrix L, composed of *m* blocks of *n × n* submatrices. In addition to the nearest-neighbor exchange elements along the θ direction, 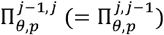 and 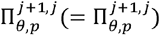, the matrix also includes nearest-neighbor off-diagonal exchange elements along the □ direction, 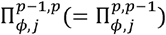 and 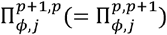. These elements are located at the matrix indices at ([*p* − 2] *n* + *j*, (*p* − 1) *n* + *j*) and ([*p* − 1] *n* + *j*, (*p* − 2) *n* + *j*) for 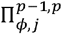, as well as ((*p* − 1) *n* + *j*, (*pn* + *j*) and (*pn* + *j*, (*p* − 1) *n* + *j*) for 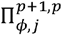, respectively, where *j* = 1,2, …, *n* and *p* = 1,2, …, *m*. The ([*p* − 1] *n* + *j*) th diagonal element of the (*nm*) × (*nm*) supermatrix consists of

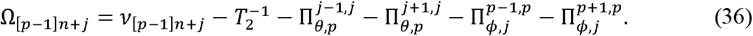

### 3.3. NMR Lineshape of Lipids within a Thinned Bilayer Dimple

Peptides and proteins interact with lipids in a bicelle, distorting its otherwise ideal geometry. In many cases the hydrophilic portion of peptides or proteins bind to the headgroups of lipids, resulting in a thinned membrane patch when this occurs within the planar region. The resulting thickness variation introduces local curvature, reorienting the bilayer normal away from that of the undisturbed flat region. Although membrane-thinning interactions can induce complex and irregular changes in bicelle structure, here we adopt a highly simplified model to qualitatively predict and analyze the structural and dynamic nature of the ^31^P, ^2^H, and ^14^N NMR lineshapes of lipids distributed on the thinned membrane surface. [38] [39] A simplified model for the thinned bilayer treats the surface as a cosine-shaped cross-section (Figure 2). Specifically, we take a single half-period of a cosine profile in the x–z plane and rotate it by 360° about the z-axis, yielding an axisymmetric dimple that mimics a locally thinned membrane patch. We model an axisymmetric dimple centered on the z-axis, characterized by a rim radius a (its maximum lateral extent) and a central depth *d* (the maximum thinning). The curved surface is characterized by a cosine cross-section,

**Figure 2.**
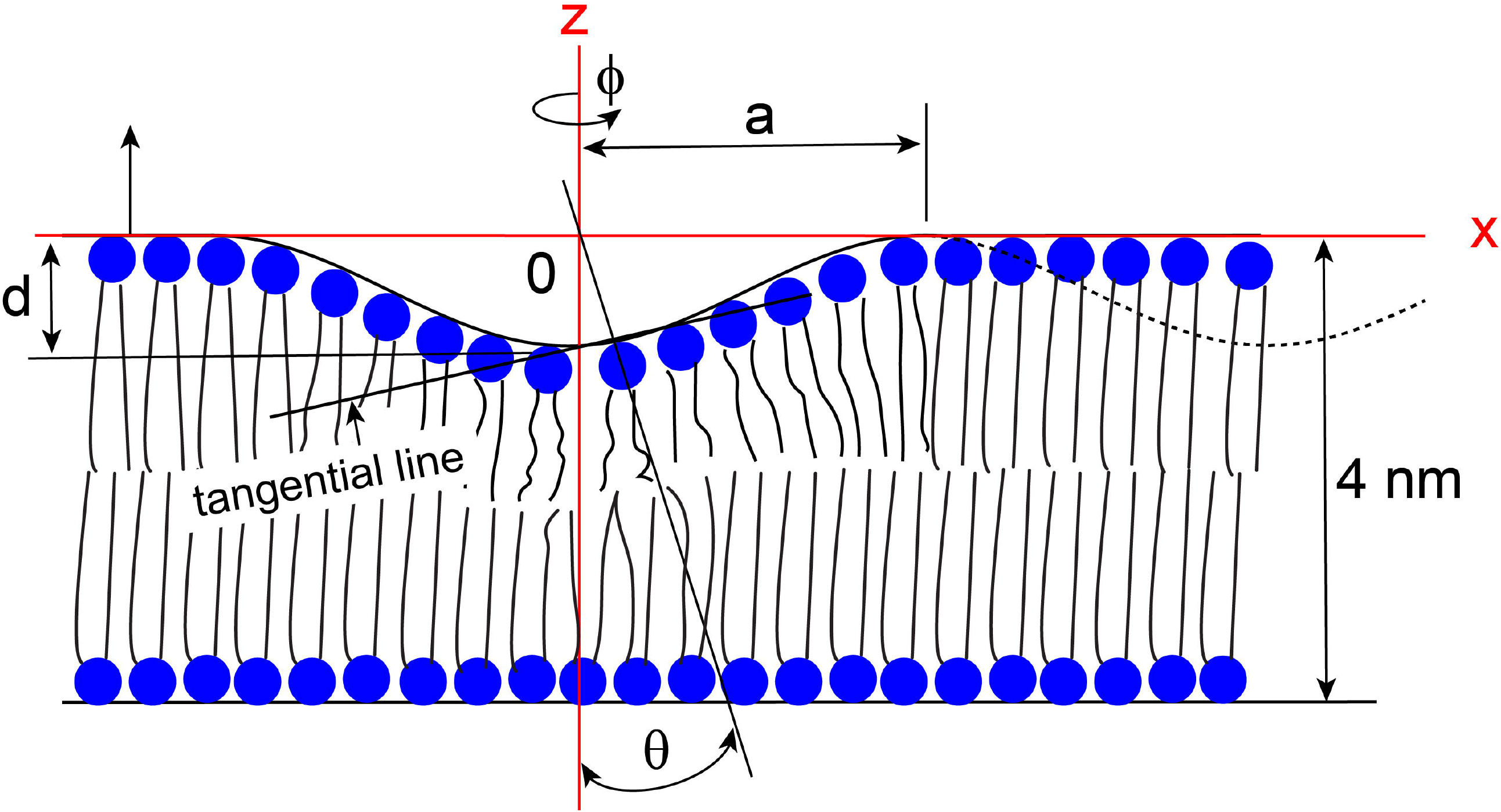
Simplified cartoon illustration of a locally thinned bilayer region induced by peptide or protein interaction with lipids. The thinned region is approximated as a concave dimple formed by rotating one period of a cosine curve, defined in the *x–z* plane, about the *z*-axis by the azimuthal angle ϕ. The depth and radius of the dimple are denoted by *d* and *a*, respectively. Lipid molecules are assumed to align perpendicular to the local membrane surface, i.e., orthogonal to the tangent of the curved membrane. The extent of membrane thinning is characterized by the ratio *d/a*. We assume the thickness of the undisturbed bilayer to be approximately ^4^ nm.

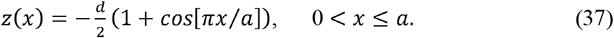

For the NMR lineshape weighting we use surface element (Jacobian), given by the product of the distance scale factors on the *xz* and *xy* planes. At fixed ϕ =0 (i.e., varying x along the meridian) we have

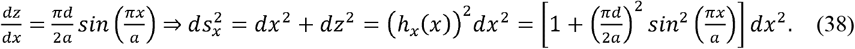

At fixed x (i.e., varying the azimuth *ϕ*,around the z), revolving gives

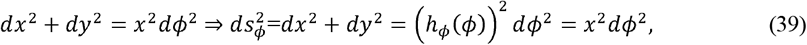

so the scale factors are

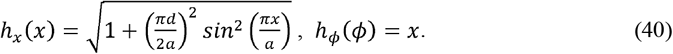

Therefore, the NMR lineshape factor (surface-area weight) at radius x (0 ≤*x* ≥*a*) is

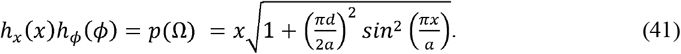

The fast uniaxial rotations of lipids along their chain axes makes a lipid align orthogonal to the tangential line drawn on the dimple surface where the lipid under consideration is positioned. When an axis is drawn that is orthogonal to the tangential line taken at the position where a lipid is located on the surface, the angle θ formed between this axis and the z-axis is provided as

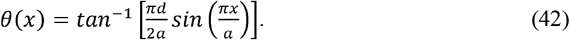

Therefore the NMR lineshape factor can be rewritten in terms of a, d, and θ by

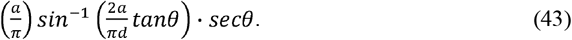

## 4. Numerical simulations of dynamic ^14^N and ^31^P NMR lineshapes of Bicelles

By explicitly accounting for both the geometrical NMR lineshape factor and lateral diffusive dynamics, experimental ^1^□N (or ^2^H) quadrupolar and ^31^P chemical shift anisotropy (CSA) NMR lineshapes arising from lipids forming a supramolecular bicelle geometry can be quantitatively simulated. Within this framework, the simulated spectra depend on several key parameters, including the quadrupole coupling or CSA tensor parameters, the bicelle q-factor, and the lateral diffusion coefficient *D*_Id_. Beyond these primary factors, the present model is sufficiently general to incorporate structural deviations from the ideal bicelle geometry. In particular, it allows numerical simulation of an elliptically distorted rim region as well as bilayer thinning in the planar disc region of bicelles. Such structural heterogeneities may naturally arise from dynamic interactions between lipid bilayers of bicelles and embedded cholesterol or peptide molecules, which can locally perturb both curvature and bilayer thickness. In the following subsections, we systematically demonstrate how each of these parameters—tensor properties, q-factor, lateral diffusion, and geometric distortions—individually and collectively influences the resulting bicelle NMR lineshapes. By varying these factors one at a time, we illustrate their distinct spectral signatures and provide a mechanistic framework for interpreting experimentally observed ^31^P and ^14^N (or ^2^H) NMR spectra of bicelles.

### 4.1. Influence of lateral lipid diffusion on bicelle NMR lineshapes

Figure 3 shows simulated dynamic ^1^□N (A, B) and ^31^P (C, D) NMR spectra of a lipid bicelle with q = 2, illustrating the dependence of the observed lineshapes on the relative lateral diffusion rate of lipids on the bicelle surface. The relative diffusion rate was systematically varied from 1, 10□^1^, 10□^2^, 10□^3^, 10□□, to 10□□, as indicated from top to bottom in each panel. Because lipid molecules undergo very fast uniaxial rotation about the acyl chain axis, which is aligned parallel to the bilayer normal, we assume axially symmetric tensor parameters (η = 0) for both the ^1^□N quadrupolar interaction and the ^31^P chemical shift anisotropy (CSA). Accordingly, the quadrupolar coupling constant and CSA span were set arbitrarily to 15 kHz for ^1^□N and 80 ppm for ^31^P, respectively, at a magnetic field strength of 9.4 T, corresponding to the Larmor frequencies of 28.9 MHz for ^1^□N and 162 MHz for ^31^P nuclei. Panels (A) and (C) correspond to bicelles with n ∥ B□, while panels (B) and (D) represent the n ⍰ B□ orientation. For each orientation, the contribution from DHPC molecules present in the rim is shown separately in red, whereas the combined spectrum including both the rim and the DMPC planar disc region is shown in black. The rim geometry was modeled with an elliptically distorted cross-section (d/b = 0.9), reflecting deviations from an ideal circular rim that may arise from lipid packing constraints or interactions with other molecules.

**Figure 3.**
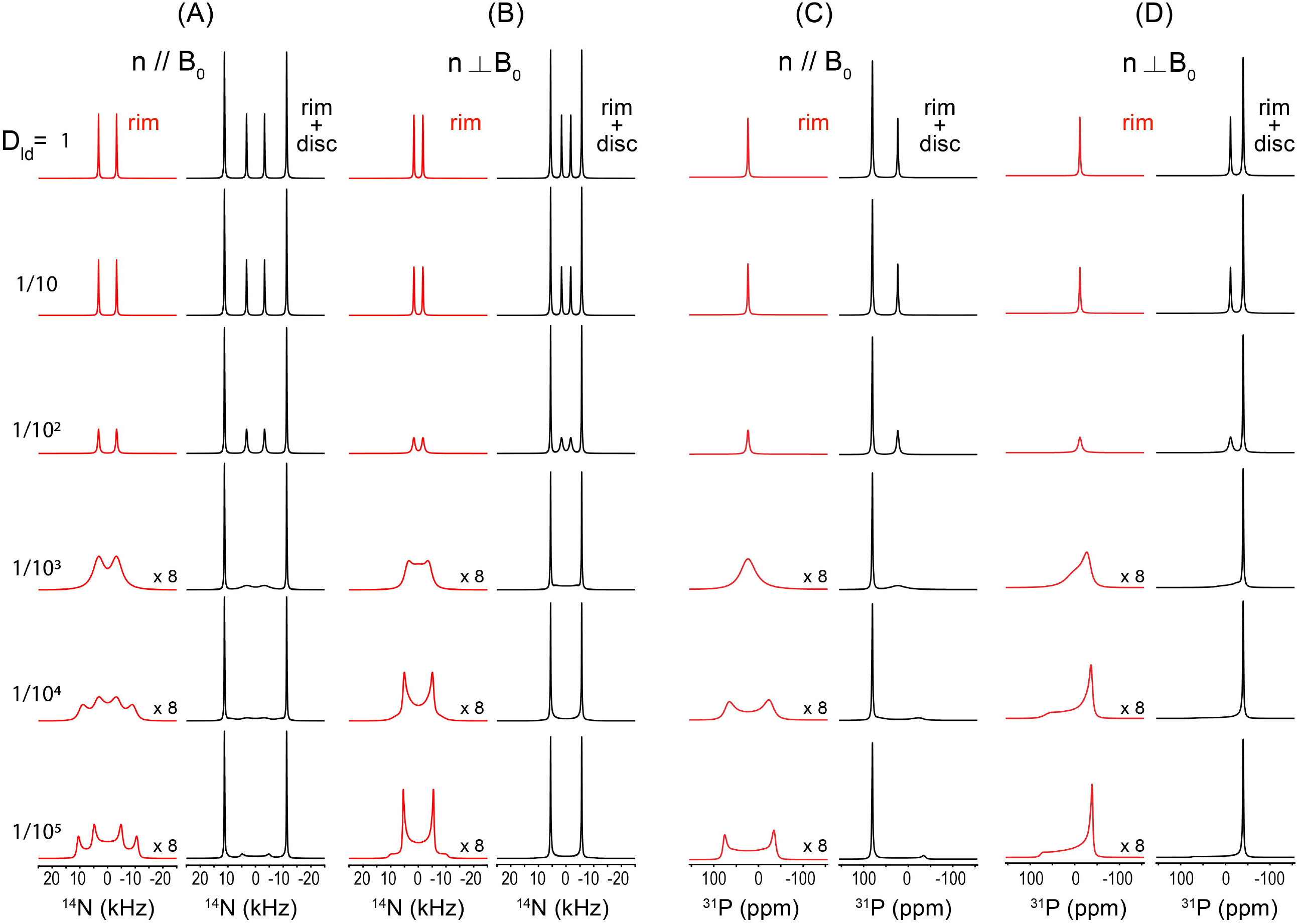
Simulated dynamic ^1^□N quadrupolar coupling and ^31^P CSA NMR spectra of a supramolecular lipid bicelles with q = 2. The simulations were performed using a ^1^□N quadrupolar coupling constant of 15 kHz (η = 0) and a ^31^P CSA of 80 ppm (η = 0) at 9.^4^ T. The rim region was modeled with an elliptical geometry characterized by d/b = 0.9, where 2b = ^4^ nm corresponds to the DMPC bilayer thickness. Panels (A) and (B) show ^1^□N spectra for bicelles with n ∥ B□ and n⍰ B□, respectively, while panels (C) and (D) present the corresponding ^31^P spectra under the same orientational conditions. For each panel, spectra are shown for progressively decreasing relative lateral diffusion rates, from D_ld_ = 1 (top) to 1/10□ (bottom). In each case, the rim contribution (DHPC region) is shown in red, whereas the combined spectrum including both the rim and the flat disc region (DMPC region) is shown in black to the right. At fast diffusion rates, the contribution from molecules present in the rim is motionally averaged into a narrow, nearly isotropic line. As the lateral diffusion rate decreases, this motional averaging becomes incomplete, and the rim spectral lines progressively broaden into anisotropic lineshapes reflecting the underlying bicelle geometry and lipid distribution. In contrast, the spectral shape from the flat disc region remains invariant in every case, as all lipids in this region possess an identical orientation with respect to the external magnetic field B□. Two hundred θorientations were used for n // B□, and 80 (θ, ϕ) orientations for n⍰ B□; all spectra were processed with 120 Hz line broadening.

At fast lateral diffusion rates, the rim-associated spectral lines are strongly motionally averaged, resulting in narrow, nearly isotropic lineshapes. As the diffusion rate decreases, this averaging becomes progressively incomplete, and the rim contribution broadens into anisotropic lineshapes that directly reflect the underlying bicelle geometry and spatial lipid distribution. This transition is clearly visible for both ^1^□N and ^31^P nuclei and for both bicelle orientations relative to B□. In contrast, the spectral contribution from the flat disc region remains essentially invariant with respect to diffusion rate and geometric anisotropy. This behavior arises because all lipids in the planar bilayer region share an identical orientation relative to B□, and lateral diffusion within this region does not introduce additional orientational averaging. One limitation of the present treatment is that the actual size of the bicelle and the corresponding number of lipid molecules are not explicitly taken into account. As a result, the diffusion coefficients discussed here should be regarded as relative rather than absolute values. In a later section, we will address this limitation by incorporating realistic bicelle sizes and physically plausible numbers of lipid molecules, enabling the extraction of absolute diffusion rates from experimentally measured ^1^□N and ^31^P NMR spectra (vide infra).

### 4.2. Spectral changes induced by variations in tensor parameters, q-value, and d/b ratio

In this case, we focus on the situation where the lipid diffusion is sufficiently fast that complete motional averaging is achieved, corresponding to the topmost row spectra in Fig. 3. Under this fully motionally averaged condition, we examine how variations in the ^1^□N quadrupolar and ^31^P CSA tensor parameters, the q-factor, and the elliptical cross-sectional shape of the rim region influence the resulting ^1^□N and ^31^P NMR spectra.

Figure 4 presents simulated NMR spectra illustrating the effects of varying the ^1^□N Cq and the ^31^P CSA. Panels (A) and (B) show the ^1^□N spectra obtained by increasing Cq from 5, 10, 15, 20, 25, to 30 kHz from top to bottom, while panels (C) and (D) display the ^31^P spectra corresponding to CSA span of 20, 40, 60, 80, 100, and 120 ppm, respectively. Assuming that DHPC and DMPC share identical CSA and QC tensor parameters, ssNMR simulations were performed for each nucleus under both n ∥ B□ (A, C) and n ⍰ B□ (B, D) orientations for the ^31^P and ^1^□N nuclei. All other parameters were held constant at the same values used in Fig. 3, namely q = 2 and d/b = 0.9, and a line-broadening factor of 120 Hz was applied uniformly to all spectra. The simulated spectra are straight-forward to interpret as the less intense peaks correspond to the molecules present in the rim and the highest intensity peaks are from lipids present in the planar lipid bilayer. As evident from the simulated results in Fig. 4, the present bicelle model faithfully produces the expected NMR lineshapes in response to systematic changes in the Cq and CSA tensor parameters, demonstrating its capability to capture the tensor-dependent spectral characteristics of bicellar lipid systems.

**Figure 4.**
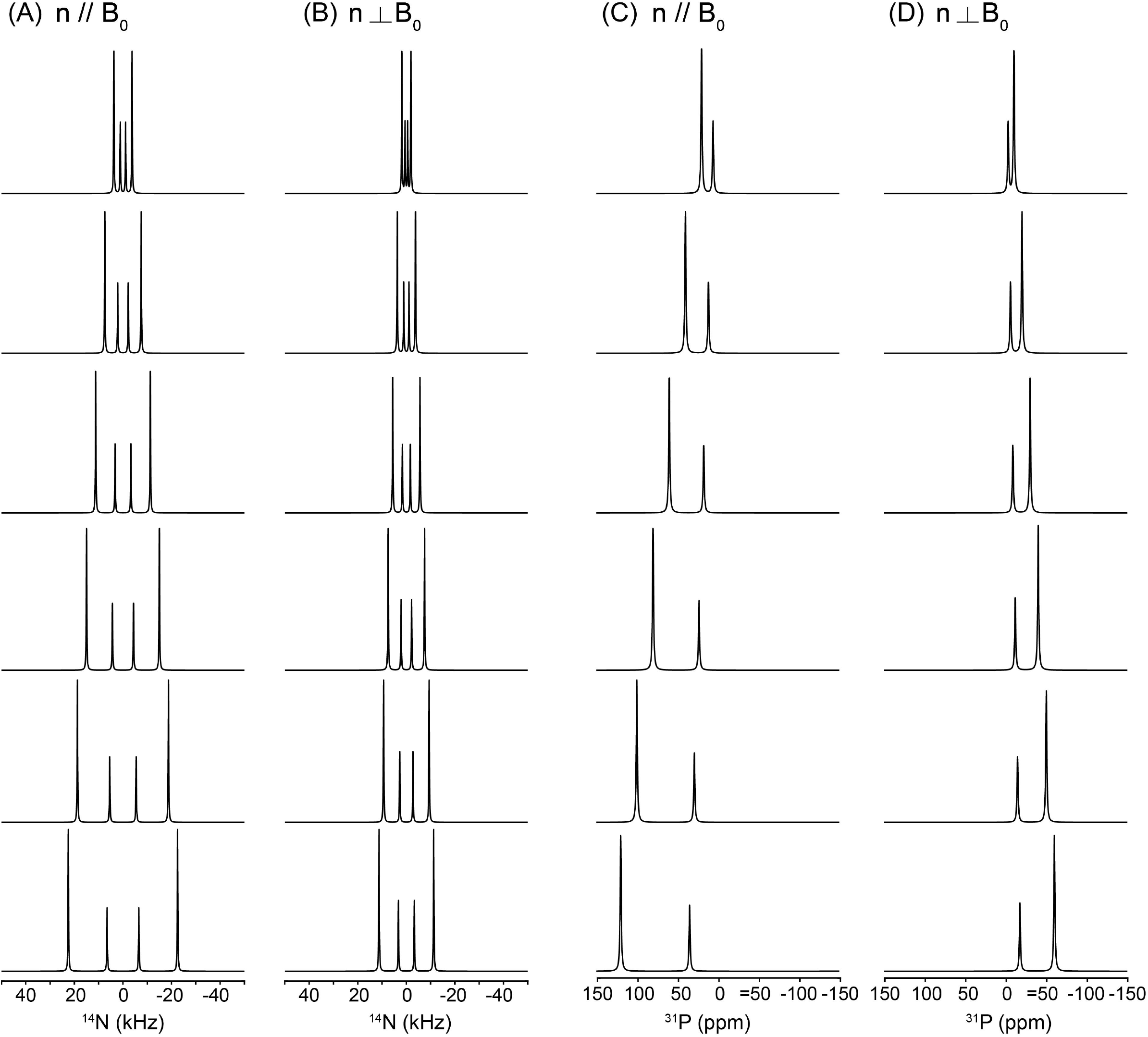
Simulated ^1^□N quadrupolar and ^31^P CSA NMR spectra of a bicelle under fully motionally averaged conditions. Panels (A) and (B) are ^1^□N NMR spectra simulated for bicelles with the director n ∥ B□ and n⍰ B□, respectively, while panels (C) and (D) present the corresponding ^31^P NMR spectra for the same two orientations. From top to bottom, the ^1^□N Cq is increased from 5, 10, 15, 20, 25, to 30 kHz in panels (A) and (B), and the ^31^P CSA span is increased from 20, ^4^0, 60, 80, 100, to 120 ppm in panels (C) and (D). All simulations were performed with q = 2 and an elliptically distorted rim geometry characterized by d/b = 0.9, identical to the parameters used in Fig. 3. A uniform line-broadening factor of 120 Hz was applied to all spectra.

Figure 5 illustrates the influence of the q-factor on the simulated ^1^□N (A, B) and ^31^P (C, D) NMR spectra of bicelles, where the q-value is varied from 0.5 to 3.5 from top to bottom. Simulations are shown for both bicelle orientations, n ∥ B□ (A, C) and n ⍰ B□ (B, D). All other parameters were held constant, with Cq = 15 kHz for ^1^□N, CSA = 80 ppm for ^31^P at 9.4 T, and a rim geometry characterized by d/b = 1, under conditions of sufficiently fast lateral diffusion leading to complete motional averaging, as in Fig. 4. Because the q-factor reflects the molar ratio of DMPC to DHPC, changes in q directly modify the relative population of molecules residing in the planar bilayer (DMPC) versus the rim (DHPC) regions. For smaller q-values, corresponding to a larger fraction of rim lipids, the inner pair of peaks in the ^1^□N quadrupolar spectra becomes more intense, whereas increasing the q-value enhances the outer pair of peaks, reflecting the growing contribution from the planar bilayer region. A similar trend is observed in the ^31^P CSA spectra, although the effect manifests as an asymmetric redistribution of peak intensities. For smaller q-values, the right-hand peak is enhanced in the n ∥ B□ orientation, while the left-hand peak becomes more intense for n ⍰ B□. As the q-value increases, these intensity patterns are reversed, consistent with the changing balance between rim and planar lipid populations. Overall, Fig. 5 demonstrates that the present bicelle model sensitively captures how variations in the DMPC/DHPC molar ratio translate into systematic and orientation-dependent changes in both ^1^□N and ^31^P NMR lineshapes, providing a direct spectroscopic handle for probing bicelle composition. It should be noted that small bicelles (i.e., those with low *q* values) do not magnetically align and therefore exhibit isotropic NMR spectra as they randomly tumble fast on the NMR time scale to motionally average out the anisotropic interactions. However, for the purpose of demonstrating the utility of the simulation framework, we assume that bicelles with all *q* values considered in this study are aligned in an external magnetic field.

**Figure 5.**
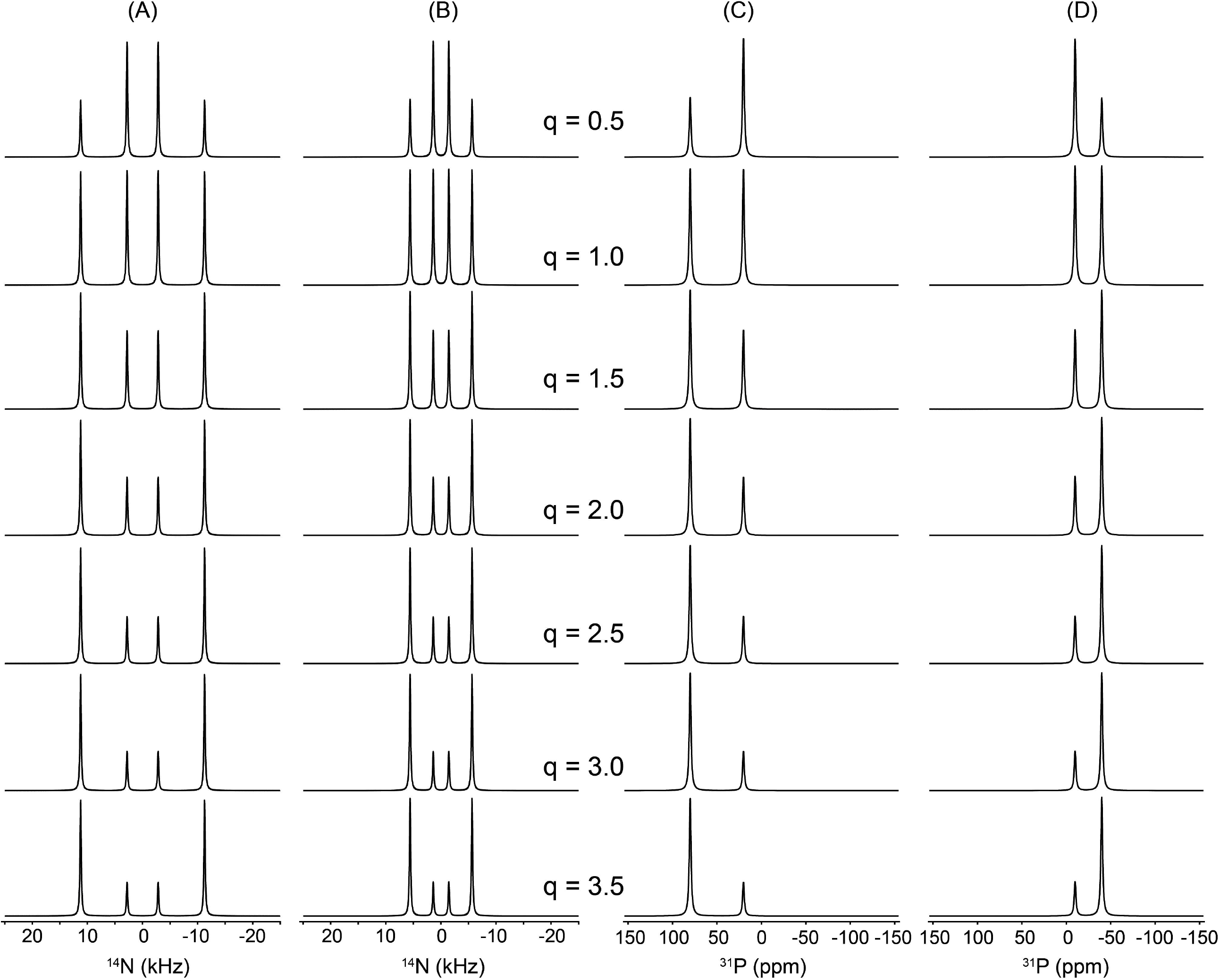
Simulated ^1^□N (A, B) and ^31^P (C, D) NMR spectra showing the effect of the q-factor (DMPC/DHPC molar ratio), varied from 0.5 to 3.5 from top to bottom, under fully motionally averaged conditions. Spectra are shown for n ∥ B□ (A, C) and n⍰ B□ (B, D). All other parameters were fixed: Cq = 15 kHz, CSA = 80 ppm at 9.^4^ T, d/b = 1, line broadening 120 Hz.

Finally, we examine the effect of geometrical deformation of the bicelle rim region, which may arise when a bicelle interacts with peptides or proteins, or due to thermal energy at high T rendering collision and fusion of the discs, leading to distortions in the rim cross section. Within our model, such geometry changes are described by variations in the ellipticity of the rim cross section, parameterized by the d/b ratio. Changes in this parameter represent deviations from an ideal circular rim and are therefore expected to have a pronounced impact on the resulting NMR lineshapes. As will be discussed later (vide infra), this variable is far from trivial and becomes critically important when simulating experimentally measured spectra. Figure 6 illustrates the influence of the d/b ratio on the simulated ^1^□N quadrupolar (A, B) and ^31^P CSA (C, D) NMR spectra of bicelles. The d/b value is varied from 1.3 to 0.7 from top to bottom. Simulations are shown for both bicelle orientations, n ∥ B□ (A, C) and n ⍰ B□ (B, D). All other parameters were fixed at Cq = 15 kHz for ^1^□N and CSA = 80 ppm for ^31^P at 9.4 T with an applied line-broadening of 120 Hz. The simulations were carried out under conditions of sufficiently fast lateral diffusion, resulting in complete motional averaging, as in Figs. 3 and 4.

**Figure 6.**
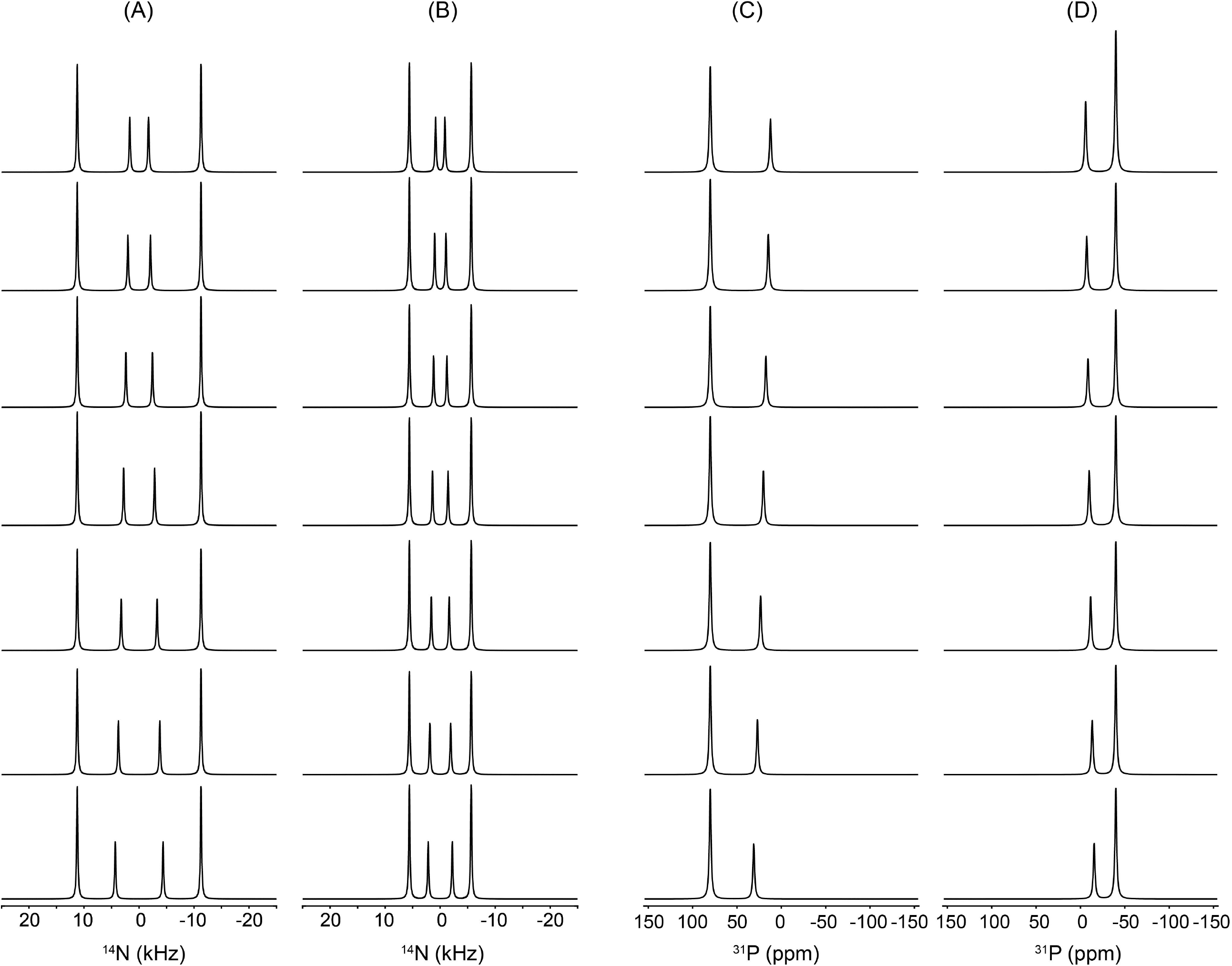
Simulated ^1^□N (A, B) and ^31^P (C, D) NMR spectra showing the effect of rim ellipticity (d/b) under fully motionally averaged conditions. The d/b ratio is varied from 1.3 to 0.7 from top to bottom for n ∥ B□ (A, C) and n⍰ B□ (B, D). All other parameters were fixed (q = 2, Cq = 15 kHz, CSA = 80 ppm at 9.^4^ T, line broadening 120 Hz). Variations in d/b affect only the rim contribution, producing systematic changes in peak separation (^1^□N) and position shifts (^31^P). This model can potentially describe the rim deformation associated with bicelle–biomolecule interactions.

The simulated spectra are easy to interpret as the variations in the d/b ratio affect exclusively the rim contribution, while the spectral features arising from the planar bilayer region remain unchanged. As the d/b value increases, the separation between the inner doublet peaks in the ^14^N spectrum becomes narrower, whereas decreasing the d/b value results in a wider separation. A similar trend is observed in the ^31^P spectra; however, because the rim contribution consists of a single spectral component, the effect manifests primarily as a systematic frequency shift. Specifically, for the n ∥ B□ orientation, increasing the d/b value shifts the rim resonance upfield (to the right), while for n ⍰ B□, it shifts to the left. The opposite behavior is observed as the d/b value decreases. Taken together, Fig. 6 demonstrates that rim ellipticity is a key geometrical parameter governing bicelle NMR lineshapes. Its influence is comparable in importance to tensor parameters and q-factor variations, underscoring the necessity of explicitly accounting for rim geometry when interpreting experimental ^1^□N and ^31^P bicelle NMR spectra, particularly in systems perturbed by peptide or protein binding.

One physically plausible validation of the rim-geometry interpretation based on a variable *d/b* ratio is that it naturally reflects rim deformations induced by orientationally or directionally selective biomolecular binding. For larger *d/b* values (*d/b* > 1, oblate form), the deformation is consistent with peptide or protein binding preferentially along an oblate direction in the *xz* plane (Fig. 1A), rather than strictly along the *x* axis when viewed in the *xz* cross section. Such binding is expected to compress the rim region, leading to an elongation of the rim cross section along the *d* axis and, consequently, to an elliptical rim geometry in which *d* becomes the major axis. Simulations of experimental ^1^□N and ^31^P ssNMR spectra under peptide-binding conditions presented in Section 5 (vide infra) provide representative examples of this scenario. Conversely, smaller *d/b* values (*d/b* < 1) may correspond to peptide binding modes oriented predominantly along the *x* axis, for which an effective reduction of *d* results in a rim cross section where the half bilayer thickness *b* becomes the major axis. In this sense, *d/b* < 1 can be regarded as a phenomenological descriptor of enhanced rim compression, without implying a unique or exclusive peptide-binding orientation.

### 4.3 From Bicelle Geometry to Realistic Lipid Populations: Implications for Dynamic ^1^_D_N and ^31^P NMR

The calculations based on the model shown above do not explicitly take into account the actual number of lipid molecules contained within a single bicelle unit. Instead, an arbitrary number of lipid molecules was included in the simulations. As a result, while the spectral lineshapes governed by the geometry factor still validly reflect realistic features of the system, the lateral diffusion rates assigned to the lipid molecules do not represent absolute, physically meaningful values. For example, in Fig. 3, the motionally averaged ^1^□N and ^31^P NMR lineshapes calculated for each row correspond to different lateral diffusion rates. However, these diffusion rates cannot be interpreted as absolute quantities. Rather, under the condition that all other simulation parameters are held constant, they should be understood only in a relative sense, i.e., as representing ratios between different diffusion regimes. More specifically, the simulated ^1^□N and ^31^P spectra shown in the topmost row of Fig. 3 can be described as corresponding to a case in which the lateral diffusion is 100 times faster than that used in the spectra calculated for the third row from the top. Beyond such relative comparisons, no direct quantitative meaning should be ascribed to the absolute values of the diffusion rates employed in the simulations.

In this subsection, we evaluate the realistic size of a bicelle unit as a function of the q value and estimate the actual number of lipid molecules that can be accommodated in the flat disc and the curved rim regions. We then discuss how these molecular counts can be incorporated into dynamic ^1^□N and ^31^P NMR lineshape simulations of bicelles. We begin by considering the bilayer half-thickness and the q-factor. For a DMPC bilayer, the total bilayer thickness 2*b* is approximately 4 nm, yielding *b* ≈ 2nm. The q-factor is typically specified during bicelle preparation as the molar ratio of DMPC to DHPC. In practice, the q-factor is not a strictly fixed parameter; rather, it would reflect a dynamic equilibrium that depends on multiple experimental variables, including temperature, pH, and the balance between aligned and isotropic bicelle phases, etc. Despite this intrinsic variability, we assume that a well-defined, effective q-factor can be assigned based on the experimental preparation conditions. With an arbitrarily chosen value of the *d* /*b* is ratio (which can be optimized iteratively), the corresponding rim half-width determined. The surface area of the flat bilayer disc, accounting for both leaflets, is then given by

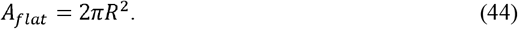

Here, *R* = *a* + *d* denotes the radius of the flat disc region. While *R* is ultimately constrained by the experimentally defined q-factor, its explicit determination requires a more detailed geometric treatment, as described below. From the geometry illustrated in Fig. 1, the surface element of the rim region can be written as

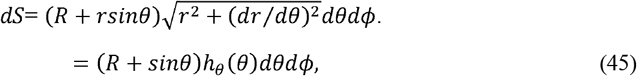

Where *h*_*θ*_ (*θ*) is the metric factor along the *θ* -direction. Integration over *ϕ* ∈ [0,2 π] and θ ∈ [0, π] yields

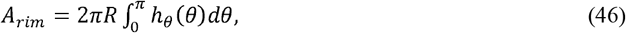

since the term 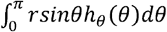 vanishes by symmetry. The integration 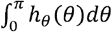 corresponds to the arc length of the rim cross-section for half a loop, i.e. approximately half of the perimeter of an ellipse. This quantity can be evaluated more conveniently by reparametrizing the rim cross-section using the ellipse parametric angle *α*, defined by *x*(*α*) = *d sin α* and z (*α*) = *b* cos *α*. With this reparameterization, Eq. (^4^6) becomes

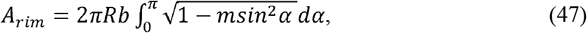

Wher 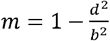 For the circular cas *A*_*rim*_ *=*2*π*^2^*Rb*. Exact evaluation of this integral requires numerical computation for an elliptic perimeter. However, for practical purposes, an accurate and simple approximation can be obtained using Ramanujan’s formul *P*_*ellipse*_ (*b,d*) for the perimeter of an ellipse, [50] [51] [52] which yields errors below approximately 0.0^4^%. Accordingly,

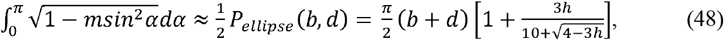

Where 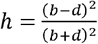. Here, the factor 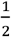 arises because only half of the elliptical perimeter considered.

The bicelle geometry can then be characterized by the ratio between the flat disc and rim regions using the q-factor, defined as [53] [5^4^]

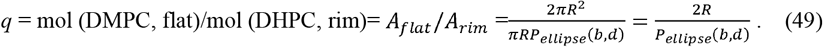

Thus, for a given q-factor and rim geometry *P*_ellipse_ (*b,d*), the radius *R* of the flat disc region can be directly determined. Here, *P*_ellipse_ (*b,d*) denotes the perimeter of the elliptical rim cross-section in the *x*– plane. To estimate the actual number of lipid molecules, we assume that each DMPC or DHPC molecule occupies an effective cross-sectional area of approximately *A*_0_ 0.62 nm^2^, projected along the bilayer normal—the axis of uniaxial lipid motion. [55] The corresponding effective molecular diameter is therefore

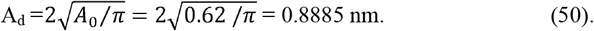

This framework enables estimation of the number of DHPC molecules in the rim region as well as the number of DMPC molecules in the flat disc region.

As an illustrative example, we consider a bicelle with q = 3.5 and d/b = 0.8. The bilayer thickness is approximately ^4^ nm, giving b = 2 nm and d = 1.6 nm. From Eq. (^4^9),

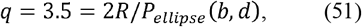

which yields

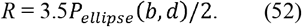

The eccentricity parameter of the elliptic rim cross section is

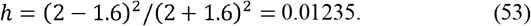

Using Ramanujan’s approximation for the ellipse perimeter,

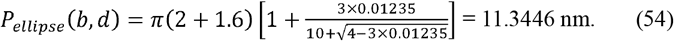

The bicelle radius R is therefore

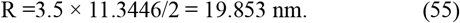

With this geometry established, the number of lipid molecules in the rim and flat disc regions can now be estimated. Using the effective molecular diameter A_d_ = 0.8885 nm obtained in Eq. (50), the number of DHPC molecules aligned along half of the rim cross-section perimeter is

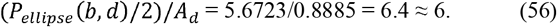

The number of DHPC molecules aligned along the outer rim perimeter at *θ*= 90° is

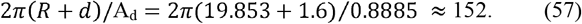

Accordingly, the total number of DHPC molecules confined to the rim surface is estimated as

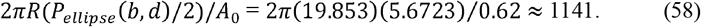

Finally, the number of DMPC molecules distributed over the flat disc region (including both leaflets) is

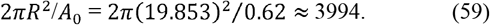

The Supporting Information provides a table summarizing lipid molecule counts calculated for several representative q-values and selected *d*/*b* ratios.

### 4.4. Lipid counts along the elliptic rim and corresponding azimuthal perimeters of the rim

Because each lipid molecule aligned along the outer half-elliptic perimeter *P*_ellipse_ (*b,d*)/2 is located at a different radial distance from the z-axis, rotation about the z-axis in the azimuthal (*ϕ*) direction generates circles with different diameters. Consequently, the number of lipid molecules that can be accommodated along these circles also varies depending on the molecular position along the rim elliptic perimeter. For highly accurate dynamic simulations of the NMR spectra, this positional dependence may also need to be taken into account. At *x* = 0, rotating the rim point at *θ* = 0° about the z-axis yields a circle of circumference 2 *πR*; rotating the rim point at *θ* = *π*/2 yields a circle of circumference 2*π*(*R* + *d*) as considered previously. For an arbitrary point *P*_ellipse_ (*x*_1_,*z*_1_), the circle formed by rotating about the z-axis has circumference 2*π*(*R* + *x*_1_). Obtaining *x*_1_ at a specified arclength position, however, necessitates a more detailed calculation. At an arbitrary point *P*_ellipse_ (*x*_1_,*z*_1_) on the elliptical perimeter, the azimuthal circle is obtained by first determining the parameter *θ*_1_ correspond to *P*_ellipse_ (*x*_1_,*z*_1_). The arclength from the the reference point to *θ*_1_ is [56] [57]

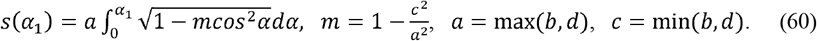

Then, the inversion *s* → *α*_1_ approximated to the second order in *m* is

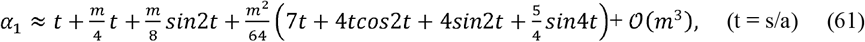

if b > d (*a = b*) and

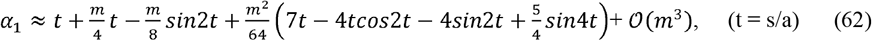

if d > b (*d = a*). Then, the coordinate of the *P*_ellipse_ (*x*_1_,*z*_1_) point is (*x*_1_,*z*_1_) = (*dsinα*_1_, *bcosα*_1_)

As an example, consider a bicelle with parameters of *q* = 3.5, *b* = 2 nm, *d* = 1.8 nm (d/b = 0.9). The perimeter of the elliptic cross-section in the *x-z* plane is calculated as

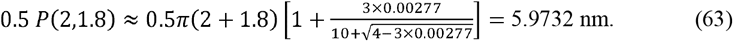

Thus, the bicelle radius is R = 3.5 × 5.9732 = 20.9062 nm. Accordingly, the number of DHPC molecules confined along the 1D elliptical perimeter of the rim is

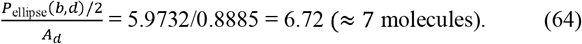

These molecules are distributed along the rim at arclengths (starting from a = 0°): 0 nm, 0.99553 nm, 1.9911 nm, 2.9866 nm, 3.9821 nm, ^4^.9777 nm, and 5.9732 nm. From above equations, these arclengths correspond to polar angles of a = 0°, 31°, 61°, 90°, 119°, 1^4^9°, and 180°, respectively. The corresponding *x*-coordinates are *xi* = R, (R + 0.9271) nm, (R + 1.57^4^3) nm, (R + 1.8) nm, (R + 1.57^4^3) nm, (R + 0.9271) nm, and R (with R = 20.9062 nm), respectively. Finally, the number of DHPC lipid molecules distributed along the azimuthal perimeter (2*πxi*) is found to be 131, 137, 1^4^1, 1^4^3, 1^4^1, 137, and 131, respectively. Table 2 summarizes these results including two other cases.

Realistic lipid packing within the bicelle is explicitly included in the dynamic ^1^□N and ^31^P NMR lineshape analyses. Figure 7 illustrates how lateral lipid diffusion on the curved rim of a q = 2 bicelle governs the static ^1^□N and ^31^P NMR lineshapes through motional averaging, while the flat disc region remains spectroscopically invariant. Simulations were performed for the two principal orientations of the bicelle normal relative to the magnetic field, n ∥ B□ and n⍰ B□, explicitly incorporating molecular counting of lipids on the elliptical rim cross-section and along the azimuthal perimeter. At low lateral diffusion coefficients (D_ld_ ≤ 10□^13^–10□^1^□ m^2^ s□^1^), the rim spectra exhibit broad, highly anisotropic powder-like patterns that reflect the wide distribution of local orientations sampled by lipids on the curved rim. These features are clearly distinguished from the disc contributions, which produce sharp, orientation-specific resonances determined solely by the fixed bilayer geometry and are independent of diffusion. As D_ld_ increases, progressive motional averaging occurs for rim lipids. The anisotropic rim patterns gradually become sharpen, and by D_ld_ ^3^ 10□^12^ m^2^ s□^1^ both the ^1^□N quadrupolar and ^31^P CSA lineshapes converge into a single, narrow resonance, characteristic of the fast-diffusion regime. This collapse reflects rapid lateral exchange among rim sites with distinct orientations, effectively averaging the anisotropic interactions over the entire rim surface. Importantly, this transition occurs for both nuclei and for both n ∥ B□ and n⍰ B□ orientations, demonstrating that the onset of the fast-diffusion regime is a robust geometric and dynamical property of the rim rather than a nucleus-specific effect. Throughout the diffusion range, the disc signals remain unchanged, underscoring that lateral diffusion primarily modulates the curved rim region while leaving the uniformly aligned flat bilayer unaffected. Together, these simulations show that lateral diffusion on the rim provides a powerful mechanism for transforming broad static powder patterns into sharp, motionally averaged resonances, and that the emergence of a single narrow rim resonance at D_ld_ ^3^ 10□^12^ m^2^ s□^1^ can be used as a direct spectral signature of fast lateral lipid mobility in bicellar assemblies. In this model, the lateral diffusion coefficient has an absolute physical meaning, as the realistic number and packing of lipid molecules are explicitly incorporated.

**Figure 7.**
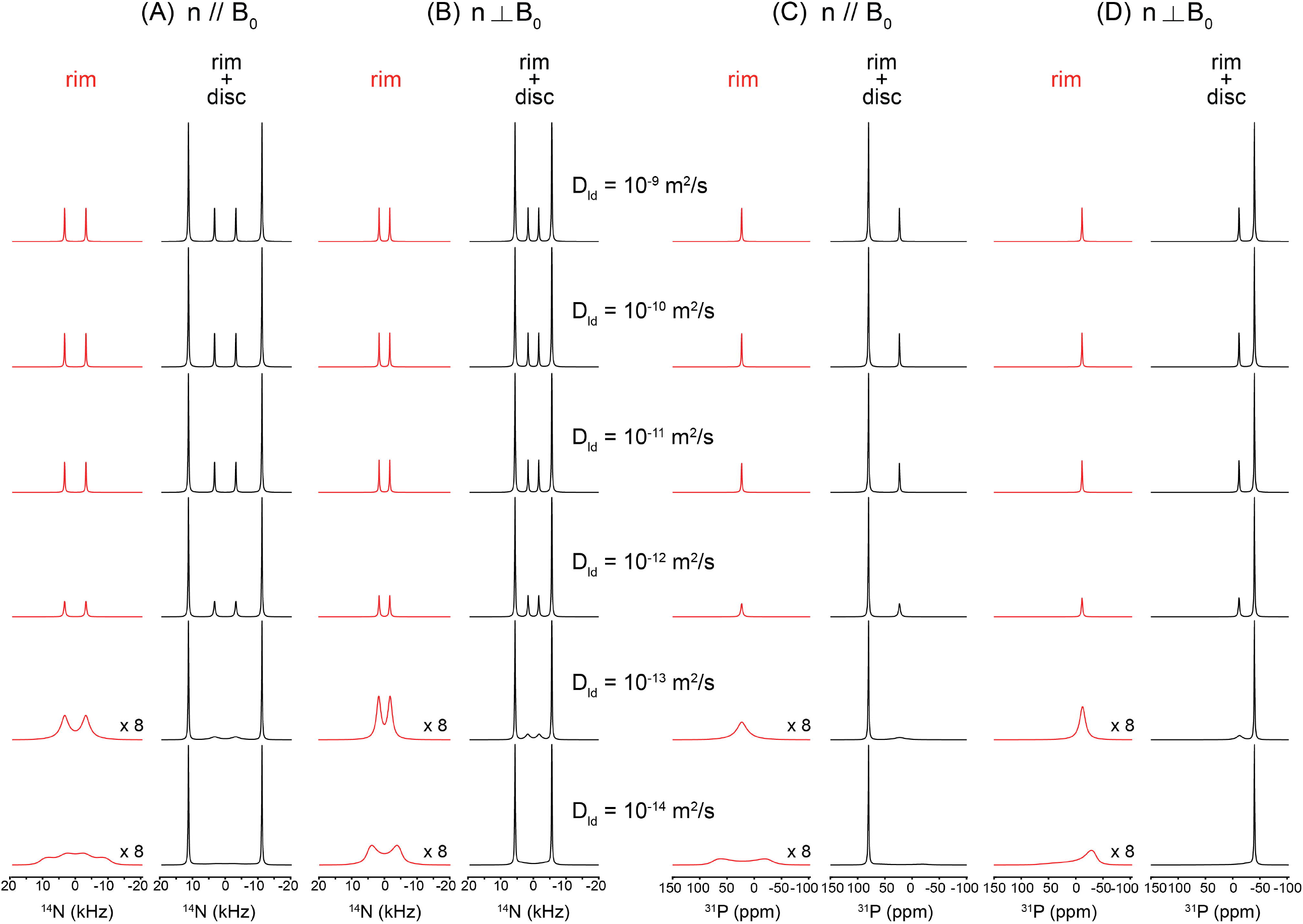
Lateral-diffusion–dependent ^1^□N and ^31^P NMR lineshape simulations for a *q* = 2 bicelle with explicit molecular counting of rim and disc lipids. Simulated static ^1^□N (left half, kHz) and ^31^P (right half, ppm) NMR lineshapes of a *q* = 2 bicelle are shown for two principal orientations of the bicelle normal relative to the external magnetic field B□: n ∥ B□ (A, C) and n ⊥ B□ (B, D). The bicelle geometry is defined by a bilayer thickness *b* = 2.0 nm (also the long axis of the elliptical rim cross-section) and a short axis *d* = 1.8 nm. Seven lipid molecules populate the outer rim cross-section, and the corresponding numbers of lipids distributed along the azimuthal perimeter at each rim position are 131, 138, 1^4^2, 1^44^, 1^4^2, 138, and 131, explicitly incorporated into the simulation (Table 1). Quadrupolar and chemical shift parameters were set to Cq (^1^□N) = 15 kHz (η = 0) and CSA(^31^P) = 80 ppm (η = 0) at the magnetic field of 9.^4^ T. For each orientation, spectra are calculated as a function of the lateral diffusion coefficient *D*_ld_ ranging from 10□□ to 10□^1^□ m^2^ s□^1^ (top to bottom). Red traces correspond to rim-only contributions, while black traces represent the combined rim + disc signals. The flat disc region, being uniformly aligned with respect to B□, yields orientation-dependent but diffusion-independent spectral patterns, whereas the rim region exhibits pronounced motional averaging as lateral diffusion increases. For both ^1^□N and ^31^P nuclei and for both bicelle orientations, the rim spectra collapse into a single motionally averaged resonance at *D*_ld_ ≥ 10□^12^ m^2^ s□^1^, marking the transition to the fast-diffusion regime. Rim spectra at the lowest diffusion coefficients *D*_ld_ ≤ 10□^13^ m^2^ s□^1^ are vertically scaled (×8) for clarity. Line broadening arising from T□ relaxation was emulated by introducing a Lorentzian line-broadening factor of 80 Hz.

### 4.5 Dynamic Spectral Lineshapes of the Flat Bicelle Disc

Changes in the structural and dynamic NMR lineshapes of lipid molecules distributed in the flat disc region occur when their orientational distribution is perturbed by the binding of peptides or proteins, such that the lipids no longer maintain a uniform orientation with respect to the external magnetic field. One example of a model that describes such a situation is the membrane thinning model discussed in the preceding subsection. [38] [39] Figure 8 illustrates the membrane-thinning impact on the dynamic ^1^□N and ^31^P NMR lineshapes of lipids in the flat disc region. The geometric distortion introduced by peptide- or protein-induced membrane thinning is represented by a simplified dimpled bilayer surface with d/a > 0 (Fig. 2), which perturbs the otherwise uniform orientational distribution of lipid molecules relative to B□. As shown in the simulated spectra in Fig. 8, the non-distorted disc lipids produce sharp, uniform-orientation resonances that are invariant with diffusion. In contrast, under slow lateral diffusion (D_ld_ ≤ 10□^1^□ m^2^ s□^1^), the lipids distributed over the distorted, thinned dimple exhibit pronounced anisotropic powder-like ^1^□N quadrupolar and ^31^P CSA lineshapes for both n ∥ B□ and n⍰ B□ orientations, reflecting the anisotropic distribution of local molecular orientations imposed by the curved dimple surface. With increasing lateral diffusion (D_ld_ ≥ 10□^13^ m^2^ s□^1^), rapid exchange among inequivalent sites leads to motional averaging of these anisotropic interactions, resulting in the collapse of the distorted lineshapes into a single averaged resonance. The resulting consequence is an apparent reduction of the effective ^1^□N quadrupolar coupling and ^31^P CSA tensor magnitudes, manifested as systematic shifts of the ^1^□N and ^31^P resonance positions toward lower frequencies as shown in Fig. 8. The extent of this shrinkage directly reflects the depth of the thinned dimple (d/a), providing a quantitative spectroscopic measure of membrane thinning under fast lateral lipid diffusion.

**Table 1.** Azimuthal perimeter–dependent distribution of DHPC lipids on the elliptic outer rim (*b* = 2.0 nm, *d* = 1.8 nm, *q* = 3.5)

| Position of lipid | $s_i$ | $s_i/b$ | $\alpha_i$ | $x_i$ | $2\pi x_i$ |
| --- | --- | --- | --- | --- | --- |
| 1 | 0 | 0 | 0 | 20.9062 | 131 |
| 2 | 0.995533 | 0.497767 | 31 | 21.9363 | 138 |
| 3 | 1.991067 | 0.995533 | 61 | 22.6554 | 142 |
| 4 | 2.9866 | 1.4933 | 90 | 22.9062 | 144 |
| 5 | 3.982133 | 1.991067 | 119 | 22.6554 | 142 |
| 6 | 4.977667 | 2.488833 | 149 | 21.9363 | 138 |
| 7 | 5.9732 | 2.9866 | 180 | 20.9062 | 131 |

( $b = 2.0$ nm, $d = 2.0$ nm, $q = 3.5$ ; circular case)
| Position of lipid | $s_i$ | $s_i/b$ | $\alpha_i$ | $x_i$ | $2\pi x_i$ |
| --- | --- | --- | --- | --- | --- |
| 1 | 0 | – | 0 | 21.9911 | 138 |
| 2 | 1.0472 | – | 30 | 22.9911 | 144 |
| 3 | 2.0944 | – | 60 | 23.7232 | 149 |
| 4 | 3.1416 | – | 90 | 23.9911 | 151 |
| 5 | 4.1888 | – | 120 | 23.7232 | 149 |
| 6 | 5.236 | – | 150 | 22.9911 | 144 |
| 7 | 6.2832 | – | 180 | 21.9911 | 138 |

( $b = 2.0$ nm, $d = 2.2$ nm, $q = 3.5$ )
| Position of lipid | $s_i$ | $s_i/b$ | $\alpha_i$ | $x_i$ | $2\pi x_i$ |
| --- | --- | --- | --- | --- | --- |
| 1 | 0 | 0 | 0 | 23.1038 | 145 |
| 2 | 0.943 | 0.42864 | 24.69 | 23.9392 | 150 |
| 3 | 1.886 | 0.85727 | 50.08 | 24.6377 | 155 |
| 4 | 2.829 | 1.28591 | 76.47 | 25.0483 | 157 |
| 5 | 3.772 | 1.71455 | 103.40 | 25.0494 | 157 |
| 6 | 4.715 | 2.14318 | 129.88 | 24.6384 | 155 |
| 7 | 5.658 | 2.57182 | 155.32 | 23.9389 | 150 |
| 8 | 6.601 | 3.00045 | 179.99 | 23.1038 | 145 |

**Figure 8.**
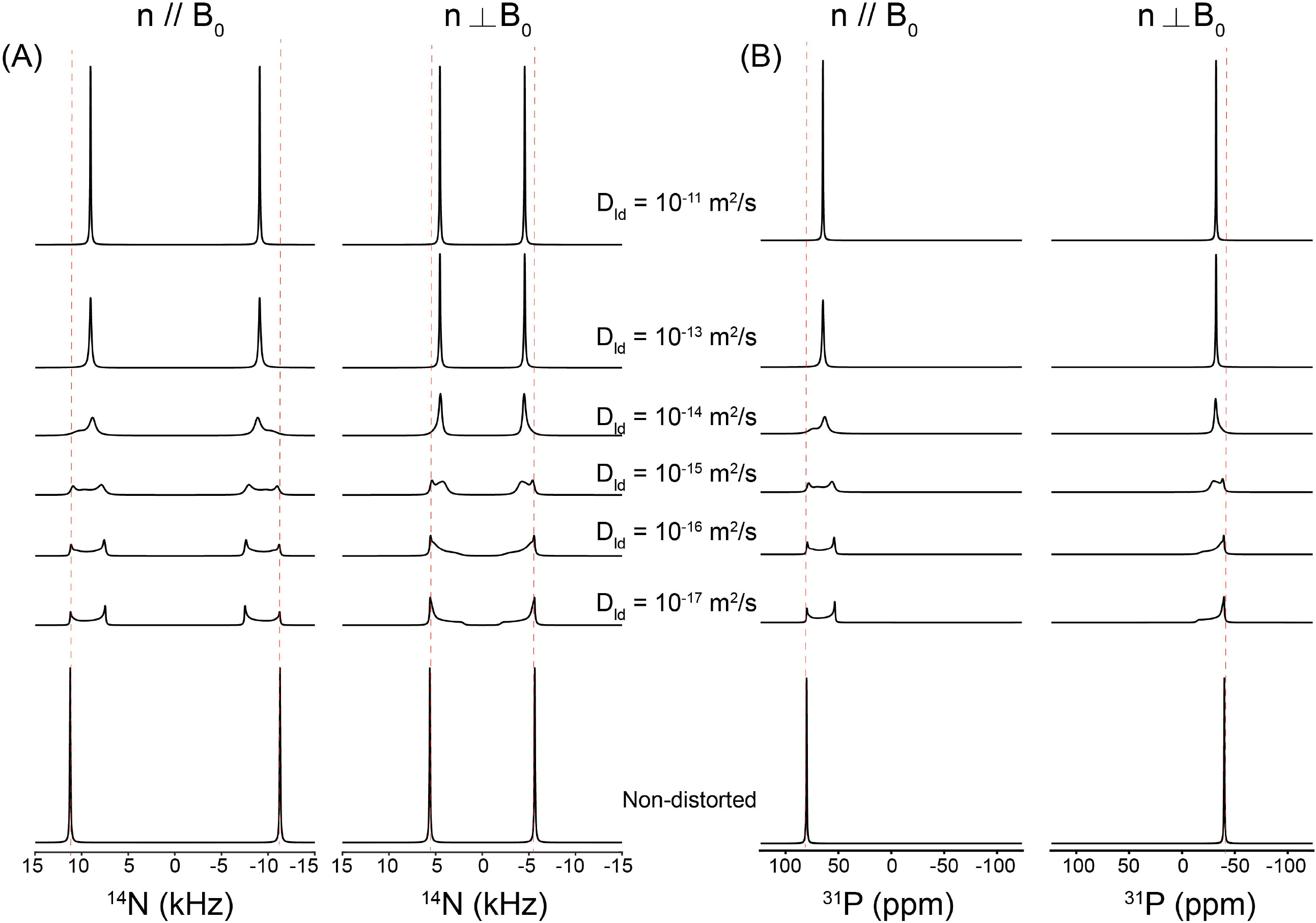
Dynamic ^1^□N and ^31^P NMR lineshape simulations for the membrane-thinning model in the flat bicelle disc region. (A) Simulated static ^1^□N quadrupolar and (B) ^31^P chemical-shift anisotropy (CSA) NMR lineshapes for lipids distributed on a hypothetical dimpled bilayer surface representing membrane thinning induced by peptide/protein binding. The thinning geometry is defined by d/a = 0.17 (see Fig. 2), with Cq(^1^□N) = 15 kHz (η = 0) and CSA(^31^P) = 80 ppm (η = 0) at 9.^4^ T. Spectra are shown for the two principal orientations of the bilayer normal relative to the magnetic field (n ∥ B□ and n⍰ B□) and as a function of the lateral diffusion coefficient D_ld_ ranging from 10□^11^ to 10□^1^□ m^2^ s□^1^ (top to bottom). The bottom traces correspond to the non-distorted, ideally flat bilayer. At slow diffusion (D_ld_ ≤ 10□^1^□ m^2^ s□^1^), distorted lipids on the thinned dimple exhibit pronounced anisotropic powder-like lineshapes due to the broad distribution of local molecular orientations. With increasing diffusion (D_ld_ ≥ 10□^13^ m^2^ s□^1^), rapid lateral exchange produces motional averaging, collapsing an anisotropic pattern into a single averaged resonance. The resulting apparent shrinkage of the effective ^1^□N quadrupolar coupling and ^31^P CSA tensors, manifested as systematic shifts of resonance positions toward lower frequencies, reflects the extent of membrane thinning specified by d/a under fast lateral diffusion. An arbitrary grid of 60 × 60 lipid orientations was employed. Because this qualitative model does not incorporate an absolute geometric scale, the lateral diffusion coefficients should be interpreted in a relative sense rather than as absolute physical rates. A Lorentzian line-broadening factor of 80 Hz was applied during data processing.

In these simulations, an arbitrary grid consisting of 60 × 60 discrete lipid orientations was employed to represent the orientational distribution of lipid molecules on the dimpled membrane surface. Because this qualitative membrane-thinning model does not explicitly incorporate an absolute geometric scale or realistic molecular packing density, the lateral diffusion coefficients introduced in the simulations should not be interpreted as absolute physical diffusion rates. Instead, the values of the lateral diffusion coefficient serve to define relative dynamical regimes—ranging from slow to fast lateral mobility—and are used to illustrate the qualitative effects of motional averaging on the resulting NMR lineshapes rather than to provide quantitative diffusion constants.

## 5. Examples of Actual Simulations of Experimentally Measured Spectra

We have now established a bicelle modeling framework that enables realistic simulations of the curved rim region of bicelles. In addition, although still qualitative in nature, we have developed the capability to simulate dynamic ^1^□N (as well as ^2^H) and ^31^P NMR lineshapes arising from thickness deformations of the flat bicelle disc region induced by intermolecular binding events. These modeling approaches were applied to experimentally acquired ^1^□N and ^31^P bicelle spectra to extract the ^1^□N quadrupolar and ^31^P CSA tensor parameters, together with the lateral diffusion rates of lipids.

Figure 9 compares experimentally measured static ^1^□N quadrupolar NMR spectra of magnetically aligned DMPC:DHPC bicelles with simulations based on the realistic bicelle model, highlighting how peptide binding and lipid composition modulate both bicelle geometry and lateral lipid dynamics. The spectrum of pure bicelles (A) exhibits the characteristic pair of sharp resonances arising from uniformly aligned lipids in the flat disc region together with motionally averaged weaker peaks from rim region (q = 3.5). Addition of Yb^3^□ ions (B) induces bicelle flipping, resulting in a wider spectral pattern consistent with reorientation of the bilayer normal parallel to B□. These two reference cases are accurately reproduced by simulations using a rim geometry with b = 2.0 nm and d = 1.^4^ nm and a lateral diffusion coefficient of D_ld_ ≥ 10□^11^ m^2^ s□^1^ (q = 3.5), validating the geometric and dynamic parameters of the model. Introduction of amphiphilic peptides produces pronounced changes in the spectral lineshapes. In the presence of desipramine (C), the experimental spectrum shows significant narrowing and displacement of the ^1^□N resonances toward lower frequencies, indicative of an apparent reduction in the effective quadrupolar coupling. Although variations in the quadrupolar coupling constant (Cq) have been reported in previous studies [58], we chose to model the observed spectral changes using a membrane-thinning framework rather than by adjusting Cq values. The experimental behavior is well reproduced by simulations that incorporate both a modified rim geometry with an increased *d/b* ratio (= 1) and a membrane-thinning deformation in the planar disc region, represented by a dimpled surface (*d/a* = 0.20) accompanied by slower lateral lipid diffusion within the thinned region. Similarly, the spectra obtained in the presence of MSI-78 (D and E) exhibit progressively slower spectral averaging and systematic shifts of the resonance frequencies toward lower values as the peptide concentration increases, consistent with enhanced membrane thinning and concomitant modifications of the rim curvature accompanied by slower lateral diffusion. These trends are quantitatively reproduced by simulations that require larger d/b ratios for the rim geometry compared to the pure bicelle case while maintaining the same Cq value, together with the inclusion of thinned disc regions characterized by slower lateral diffusion. This analysis demonstrates that MSI-78 binding induces both geometric distortion of the bicelle and significant changes in lipid mobility.

**Figure 9.**
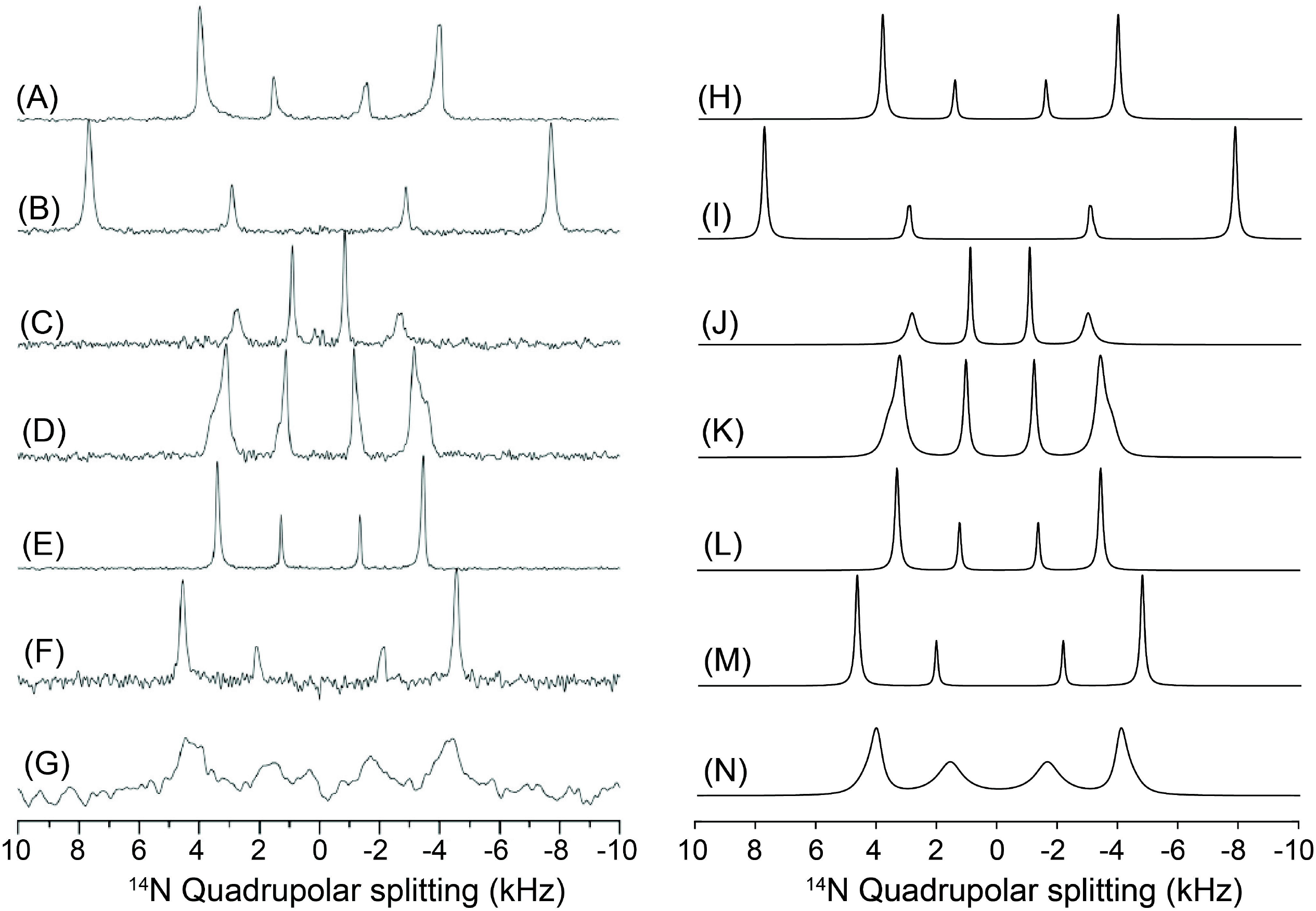
Experimental (A-G) and simulated (H-N) ^1^□N quadrupolar NMR spectra of magnetically aligned DMPC:DHPC bicelles under various perturbing conditions. Experimental ^1^□N NMR spectra of magnetically aligned DMPC:DHPC bicelles (q = 3.5) are shown for (A) pure bicelles; (B) bicelles in the presence of Yb^3^□ ions, which induce bicelle flipping; (C) DMPC:DHPC bicelles containing 2.0 mol % desipramine; (D) DMPC:DHPC bicelles containing 2.0 mol % MSI-78; (E) DMPC:DHPC bicelles containing 0.5 mol % MSI-78; (F) bicelles containing 20.0 mol % negatively charged lipid DMPG in the presence of 150 mM NaCl; and (G) the same sample in the absence of NaCl. Corresponding simulated spectra are shown in (H) and (I) for (A) and (B), respectively, for the n⍰ B□ and n ⍰ B□ orientations. Simulations used Cq = 15 kHz (η = 0), rim geometry b = 2.0 nm and d = 1.^4^ nm, and a lateral diffusion coefficient D_ld_ ≥ 10□^11^ m^2^ s□^1^ for the rim region for both cases. For (C), simulations employed Cq = 10.^4^ kHz, b = d = 2.0 nm, and D_ld_ ≥ 10□^11^ m^2^ s□^1^ for the rim region, together with a membrane-thinning model for the flat disc region characterized by d/a = 0.2 and D_ld_ = 10□^13^ m^2^ s□^1^ (q-factor = 1). For the 2.0 mol % MSI-78 sample (D), parameters were Cq = 10.^4^ kHz, b = 2.0 nm, d = 1.8 nm, and D_ld_ = 10□^12^ m^2^ s□^1^ for the rim, with a thinned disc region defined by d/a = 0.1^4^ and D_ld_ = 10□^1^□ m^2^ s□^1^ (q-factor = 2.5). For the 0.5 mol % MSI-78 sample (E), simulations used Cq = 10.^4^ kHz, b = 2.0 nm, d = 1.6 nm, and D_ld_ = 10□^11^ m^2^ s□^1^ for the rim, together with a thinned disc region characterized by d/a = 0.1^4^ and D_ld_ = 10□^11^ m^2^ s□^1^ (q-factor = 3). For the DMPG-containing bicelles (F and G), the simulations employed Cq = 12.6 kHz (η = 0) with a rim geometry defined by b = 2.0 nm and d = 1.2 nm (q = 3.5) for (F), or d = 1.6 nm (q = 1) for (G). The lateral diffusion coefficients used for the rim region were D_ld_ = 10□^11^ m^2^ s□^1^ for (F) and D_ld_ = 10□^13^ m^2^ s□^1^ for (G). In the absence of NaCl (G), an additional thinned disc region was introduced, characterized by d/a = 0.1^4^ and D_ld_ = 10□^14^ m^2^ s□^1^. An arbitrary 60 × 60 grid of lipid orientations was employed for simulations of the thinned disc region, and a Lorentzian line-broadening factor of 80 Hz was applied during data processing for all cases. Spectra A to G are reproduced from Dvinskikh et al 2006 (doi: 10.1016/j.jmr.2006.10.00^4^) with copyright permission @ Elsevier.

Bicelles containing the negatively charged lipid DMPG (F and G) exhibit furtherly pronounced spectral perturbations. In the presence of NaCl (F), the spectra remain relatively sharp but require a bigger Cq with a reduced rim diameter and faster lateral diffusion to reproduce the experimental pattern, reflecting electrostatic screening and preservation of membrane integrity. In contrast, in the absence of NaCl (G), the spectrum shows substantial broadening with slowed lateral diffusions, indicating strong perturbation of both the rim and disc regions. This feature in (G) is faithfully reproduced only when an additional thinned disc region with slow lateral diffusion is included in the simulation, consistent with enhanced electrostatic repulsion and pronounced membrane thinning.

Fig. 10 shows how binding of the antimicrobial peptide MSI-78 progressively perturbs both the curved rim and the flat disc regions of magnetically aligned DMPC:DHPC bicelles, as directly reflected in their static ^31^P NMR lineshapes. The spectrum of the pure bicelle sample (A) displays two well-resolved resonances corresponding to DHPC lipids in the rim region and DMPC lipids in the flat disc region, consistent with a bicelle geometry and rapid lateral lipid diffusion (D_ld_ ≥ 10^-10^ m^2^/s). This reference spectrum is well reproduced by simulations using an elliptic rim geometry with d/b = 0.8 (b = 2 nm; d = 1.6 nm) and no thinning deformation in the flat disc region with q = 3. Upon addition of 0.5 mol % MSI-78 (B), both resonances shift toward lower frequencies and become partially averaged, indicating perturbation of lipid orientations in both regions. These spectral changes are quantitatively captured by simulations that require an increase of the rim ellipticity to d/b = 0.9 together with the introduction of a thinned disc region characterized by d/a = 0.17, reflecting membrane thinning induced by peptide binding. At higher peptide loading (2.0 mol %, C), the frequency shifts become more pronounced to the lower frequency. Simulations reproduce these features when a nearly circular rim cross-section (d/b = 1.0) and a more strongly thinned disc region (d/a = 0.23) are introduced, demonstrating that MSI-78 binding produces both enhanced rim curvature modulation and deeper membrane thinning. Collectively, these results show that MSI-78 binding induces progressive structural deformation of bicelles while maintaining the dynamic feature in lipid lateral mobility, and that the combined rim–disc bicelle model provides a physically intuitive framework for quantitatively interpreting peptide-induced membrane perturbations from static ^31^P NMR spectra.

**Figure 10.**
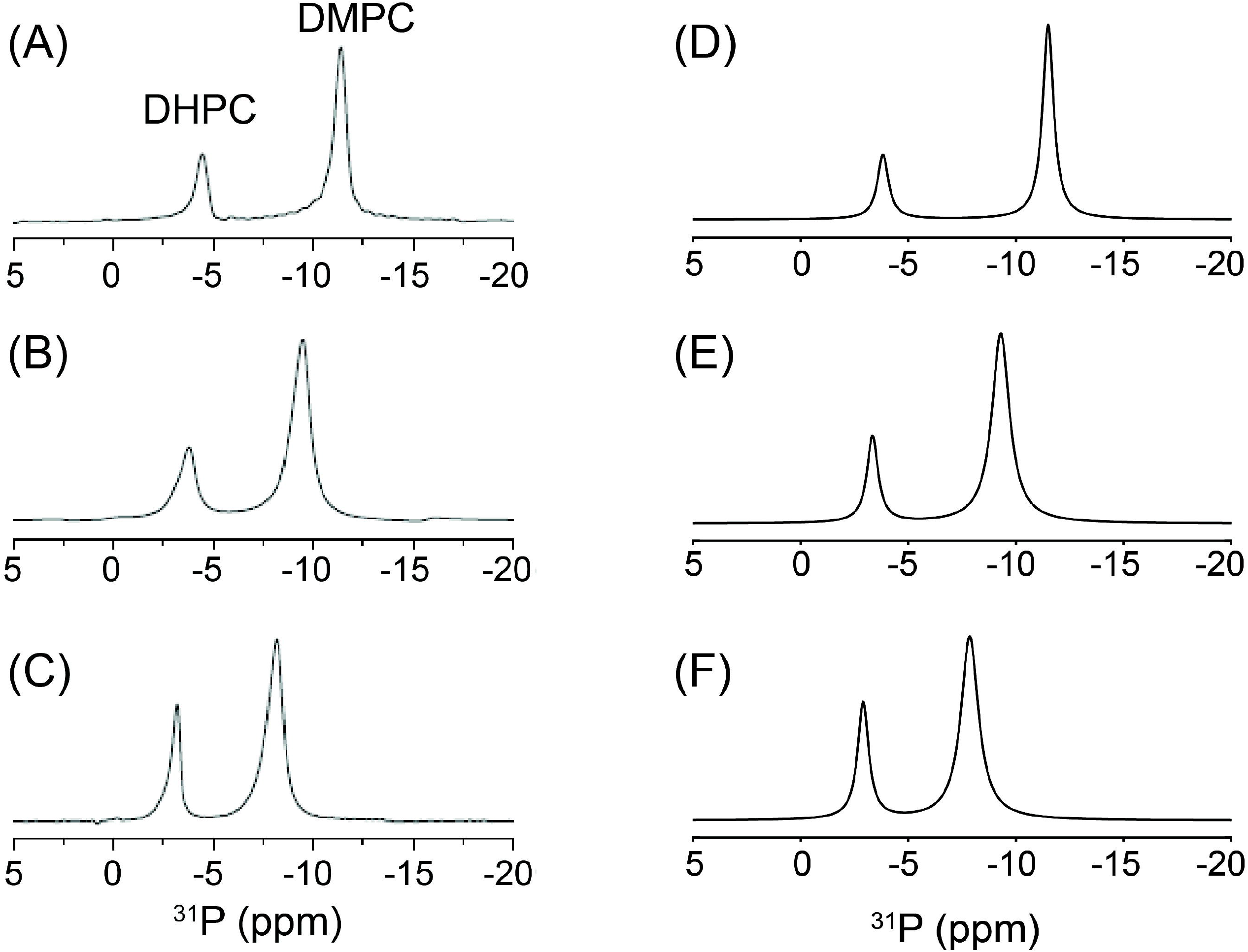
Experimental and simulated ^31^P NMR spectra of DMPC:DHPC bicelles in the presence of MSI-78. Experimental static ^31^P NMR spectra of magnetically aligned DMPC:DHPC bicelles at 9.^4^ T are shown for samples containing (A) 0 mol %, (B) 0.5 mol %, and (C) 2.0 mol % MSI-78 peptide. Spectra were acquired following a 90° radiofrequency pulse under 10 kHz ^1^H decoupling. Sixty-four transients were accumulated with a recycle delay of 3 s. The chemical-shift scale was referenced by setting the isotropic resonance observed at 10 °C to 0 ppm. Corresponding simulations of the rim region are shown in (D–F). The rim spectra were reproduced using a ^31^P CSA of 23 ppm (η = 0) with progressively increasing rim ellipticity (d/b = 0.8, 0.9, and 1.0 for A–C, respectively) and a lateral diffusion coefficient D_ld_ ≥ 10^-11^ m^2^ s□^1^. The systematic shift of the rim resonances toward lower frequency with increasing MSI-78 concentration reflects elongation of the rim cross-section along the axis orthogonal to the bilayer normal. For the flat disc region, no thinning deformation was required for the pure bicelle (A), which was simulated with q = 3. In contrast, MSI-78 binding induces progressive membrane thinning of the disc region, which was reproduced by introducing a dimpled disc surface with d/a = 0.17 (q = 2.5) for 0.5 mol % MSI-78 (B) and d/a = 0.23 (q = 2.5) for 2.0 mol % MSI-78 (C). The increasing d/a values account for the systematic downfield shifts of the disc resonances and demonstrate that MSI-78 binding produces both rim-curvature modulation and thickness deformation of the flat bicelle disc region. An arbitrary 60 × 60 grid of lipid orientations was employed for simulations of the thinned disc region, and a Lorentzian line-broadening factor of 80 Hz was applied during data processing for all cases. Spectra A to C are reproduced from Dvinskikh et al 2006 (doi: 10.1021/ja061153a) with copyright permission @ American Chemical Society.

Collectively, these simulation results demonstrate that the combined rim–disc bicelle model, together with the membrane-thinning formalism, provides a unified and physically intuitive framework for interpreting how peptide/protein binding, ionic strength, and lipid composition modulate bicelle geometry, lipid lateral mobility, and effective quadrupolar coupling tensors, as directly manifested in experimentally observed ^1^□N NMR lineshapes. These simulations demonstrate that peptide/protein binding and lipid composition modulate both the effective quadrupolar coupling and lateral lipid mobility through perturbations of the rim geometry and thickness of the flat bicelle disc region.

A controversial aspect in analyzing the flat-disc region is whether the experimentally observed reduction in apparent quadrupolar coupling (Cq) or CSA should be simulated by artificially decreasing the tensor magnitude itself or by invoking membrane-thinning dynamics. From a physical standpoint, a direct reduction of Cq or CSA is not appropriate, because peptide or protein binding creates a more crowded and sterically constrained local environment within the lipid bilayer, which would be expected to *reduce* rather than enhance motional mobility. Consequently, a smaller motionally averaged anisotropic frequency associated with peptide/protein binding cannot originate from increased molecular motion, leading to an intrinsic physical contradiction if tensor scaling is used as the fitting parameter. In contrast, invoking membrane thinning—while maintaining fast lateral lipid diffusion—provides a physically consistent mechanism that naturally reduces the effective anisotropic frequencies through geometric reorientation of lipid segments, without requiring unphysical increases in mobility. Importantly, membrane thinning induced by surface-bound peptides or proteins has been independently observed by complementary structural techniques, including neutron scattering, [59] X-ray reflectivity, [60], small angle X-ray scattering, [61] and theoretical modeling and simulations, [62] providing an experimentally verifiable and physically grounded basis for this modeling approach.

One limitation of the present model that should be noted is that lateral diffusion coefficients (D_ld_) cannot be extracted from the flat disc region. Even when the membrane-thinning model is employed, the analysis yields only relative trends among differently composed samples, rather than quantitative diffusion values. In contrast, for the bicelle rim region, where the geometry is explicitly defined and the actual number of lipid molecules contained within the geometry is realistically counted and incorporated into the simulations, the D_ld_ values extracted from the ^14^N and ^31^P NMR lineshape analyses have absolute physical meaning. Nevertheless, even in this case certain limitations remain. When the experimental peaks are already fully motionally averaged, only the slowest bound of D_ld_ can be specified, whereas faster bound of diffusion rates cannot be uniquely determined. Accordingly, such results are expressed in the form of lower limits (e.g., D_ld_ ≥ 10□^11^ m^2^ s□^1^). Despite these limitations, we believe that the ability to measure lateral lipid diffusion in the bicelle rim region using this model represents a groundbreaking example of accessing lipid dynamics at the macromolecular (mesoscopic) level.

## 3. Conclusions

Here, we introduce an elliptic bicelle geometry model for simulating experimental ^1^□N (or ^2^H) and ^31^P solid-state NMR (ssNMR) spectra of DMPC/DHPC bicelles and for quantitatively describing their structural perturbations upon binding of membrane-active peptide ligands. The exceptionally close agreement between simulated and experimental spectra provides strong validation of the proposed framework and demonstrates its suitability for extracting bicelle geometry and membrane perturbations at the molecular level. We have developed a comprehensive simulation protocol that reproduces ^1^□N (and ^2^H) quadrupolar-coupling (QC) and ^31^P chemical-shift-anisotropy (CSA) ssNMR lineshapes as explicit functions of bicelle geometry and lateral lipid diffusive dynamics. In contrast to conventional phenomenological or purely orientational-averaging approaches, the present framework incorporates explicit molecular counting, enabling quantitative determination of the number of lipid molecules participating in a bicelle at a given *q*-value and their distribution between the planar bilayer region and the rim. To the best of our knowledge, this is the first NMR lineshape simulation approach that simultaneously integrates bicelle geometry, realistic molecular population statistics, and lipid motional dynamics within a unified physical model.

Application of the model to experimental ^1^□N and ^31^P ssNMR spectra yields excellent reproduction of the observed lineshapes and allows reliable extraction of intrinsic QC and CSA tensor parameters, as well as lateral lipid diffusion rates characteristic of the bicelle system. Importantly, the extracted tensor parameters provide direct access to lipid order parameters, enabling quantitative assessment of bicelle mobility and motional averaging under varying compositional and experimental conditions. Beyond describing static bicelle structures, the model further captures ssNMR lineshape deformations arising from membrane perturbations, such as bilayer thinning induced by peptide, protein, or lipid-binding ligand association. By introducing a smoothly varying, cosine-like dimple geometry coupled with lateral lipid diffusion, the simulations reproduce experimentally observed spectral distortions in a physically intuitive and quantitatively consistent manner.

## Acknowledgements

This work was performed at the National High Magnetic Field Laboratory, which is supported by National Science Foundation Cooperative Agreement No. DMR-2128556 and the State of Florida. A.R. Acknowledges the funding support from the National Institutes of Health grant R35GM13973.

## Declaration of generative AI use

In calculating the elliptic arc length of the rim region—used to determine the lipid counts along the elliptic rim and the corresponding azimuthal perimeters—we employed AI-assisted methods to obtain the appropriate mathematical expressions. Specifically, these expressions correspond to Eqs. (^4^8)–(62).

## Notes

### Competing Interest Statement

The authors have declared no competing interest.

