## Supplemental Information for "A Dynamic NMR Lineshape Simulation Framework for Lipid Diffusion and Membrane Thinning in Bicelles and Nanodiscs"

**Supplementary Material**

1. **Bicelle Molecular Counts**

Below in the table, the 2πR^2^/A_0_ column represents the number of lipid molecules located in two disc regions, whereas 2πRP_1/2_(b,d)/A_0_ corresponds to the number of lipid molecules in the rim region. All calculated numbers must be rounded up to the nearest integer.

**Table S1. Bicelle Surface Areas and Molecular Counts as Functions of Key Parameters q-value and d/b ratio (A_0_ = 0.62 nm^2^; A_d_ = 0.8885 nm)**

| **q** | **b (nm)** | **d (nm)** | **P_1/2_(b,d) (nm)** | **R (nm)** | **P_1/2_(b,d)/A_d_** | **2πR/A_d_** | **2π(R+d)/A_d_** | **2πR^2^/A_0_** | **2πRP_1/2_(b,d)/A_0_** |
| --- | --- | --- | --- | --- | --- | --- | --- | --- | --- |
| 0.5 | 2 | 1 | 4.843594 | 2.421797 | 5.451428 | 17.12616 | 24.19784 | 59.4379 | 118.8758 |
| 0.5 | 2 | 1.2 | 5.10519 | 2.552595 | 5.745852 | 18.05112 | 26.53713 | 66.03161 | 132.0632 |
| 0.5 | 2 | 1.4 | 5.382313 | 2.691157 | 6.057753 | 19.03099 | 28.93133 | 73.39493 | 146.7899 |
| 0.5 | 2 | 1.6 | 5.672323 | 2.836162 | 6.384157 | 20.05642 | 31.3711 | 81.51735 | 163.0347 |
| 0.5 | 2 | 1.8 | 5.973159 | 2.986579 | 6.722745 | 21.12012 | 33.84914 | 90.39329 | 180.7866 |
| 0.5 | 2 | 2 | 6.283184 | 3.141592 | 7.071676 | 22.21632 | 36.35967 | 100.0202 | 200.0404 |
| 0.5 | 2 | 2.2 | 6.601083 | 3.300542 | 7.429469 | 23.34036 | 38.89805 | 110.3973 | 220.7946 |
| 0.5 | 2 | 2.4 | 6.925785 | 3.462892 | 7.794918 | 24.48845 | 41.46047 | 121.5251 | 243.0503 |
| 0.5 | 2 | 2.6 | 7.256405 | 3.628203 | 8.167029 | 25.65747 | 44.04383 | 133.4047 | 266.8094 |
| 0.5 | 2 | 2.8 | 7.592211 | 3.796106 | 8.544976 | 26.84483 | 46.64552 | 146.0376 | 292.0752 |
| 1 | 2 | 1 | 4.843594 | 4.843594 | 5.451428 | 34.25232 | 41.324 | 237.7516 | 237.7516 |
| 1 | 2 | 1.2 | 5.10519 | 5.10519 | 5.745852 | 36.10225 | 44.58826 | 264.1264 | 264.1264 |
| 1 | 2 | 1.4 | 5.382313 | 5.382313 | 6.057753 | 38.06198 | 47.96232 | 293.5797 | 293.5797 |
| 1 | 2 | 1.6 | 5.672323 | 5.672323 | 6.384157 | 40.11283 | 51.42751 | 326.0694 | 326.0694 |
| 1 | 2 | 1.8 | 5.973159 | 5.973159 | 6.722745 | 42.24024 | 54.96926 | 361.5732 | 361.5732 |
| 1 | 2 | 2 | 6.283184 | 6.283184 | 7.071676 | 44.43264 | 58.57599 | 400.0807 | 400.0807 |
| 1 | 2 | 2.2 | 6.601083 | 6.601083 | 7.429469 | 46.68072 | 62.23841 | 441.5893 | 441.5893 |
| 1 | 2 | 2.4 | 6.925785 | 6.925785 | 7.794918 | 48.97691 | 65.94893 | 486.1005 | 486.1005 |
| 1 | 2 | 2.8 | 7.592211 | 7.592211 | 8.544976 | 53.68966 | 73.49035 | 584.1503 | 584.1503 |
| 1.5 | 2 | 1 | 4.843594 | 7.26539 | 5.451428 | 51.37849 | 58.45016 | 534.9411 | 356.6274 |
| 1.5 | 2 | 1.2 | 5.10519 | 7.657785 | 5.745852 | 54.15337 | 62.63938 | 594.2845 | 396.1897 |
| 1.5 | 2 | 1.4 | 5.382313 | 8.07347 | 6.057753 | 57.09296 | 66.99331 | 660.5544 | 440.3696 |
| 1.5 | 2 | 1.6 | 5.672323 | 8.508485 | 6.384157 | 60.16925 | 71.48393 | 733.6562 | 489.1041 |
| 1.5 | 2 | 1.8 | 5.973159 | 8.959738 | 6.722745 | 63.36036 | 76.08938 | 813.5396 | 542.3598 |
| 1.5 | 2 | 2 | 6.283184 | 9.424776 | 7.071676 | 66.64896 | 80.79231 | 900.1817 | 600.1211 |
| 1.5 | 2 | 2.2 | 6.601083 | 9.901625 | 7.429469 | 70.02108 | 85.57877 | 993.5759 | 662.3839 |
| 1.5 | 2 | 2.4 | 6.925785 | 10.38868 | 7.794918 | 73.46536 | 90.43738 | 1093.726 | 729.1508 |
| 1.5 | 2 | 2.6 | 7.256405 | 10.88461 | 8.167029 | 76.97242 | 95.35878 | 1200.642 | 800.4283 |
| 1.5 | 2 | 2.8 | 7.592211 | 11.38832 | 8.544976 | 80.53448 | 100.3352 | 1314.338 | 876.2255 |
| 2 | 2 | 1 | 4.843594 | 9.687187 | 5.451428 | 68.50465 | 75.57632 | 951.0064 | 475.5032 |
| 2 | 2 | 1.2 | 5.10519 | 10.21038 | 5.745852 | 72.20449 | 80.69051 | 1056.506 | 528.2529 |
| 2 | 2 | 1.4 | 5.382313 | 10.76463 | 6.057753 | 76.12395 | 86.0243 | 1174.319 | 587.1594 |
| 2 | 2 | 1.6 | 5.672323 | 11.34465 | 6.384157 | 80.22566 | 91.54035 | 1304.278 | 652.1388 |
| 2 | 2 | 1.8 | 5.973159 | 11.94632 | 6.722745 | 84.48048 | 97.2095 | 1446.293 | 723.1463 |
| 2 | 2 | 2 | 6.283184 | 12.56637 | 7.071676 | 88.86528 | 103.0086 | 1600.323 | 800.1615 |
| 2 | 2 | 2.2 | 6.601083 | 13.20217 | 7.429469 | 93.36144 | 108.9191 | 1766.357 | 883.1786 |
| 2 | 2 | 2.4 | 6.925785 | 13.85157 | 7.794918 | 97.95381 | 114.9258 | 1944.402 | 972.201 |
| 2 | 2 | 2.6 | 7.256405 | 14.51281 | 8.167029 | 102.6299 | 121.0163 | 2134.475 | 1067.238 |
| 2 | 2 | 2.8 | 7.592211 | 15.18442 | 8.544976 | 107.3793 | 127.18 | 2336.601 | 1168.301 |
| 2.5 | 2 | 1 | 4.843594 | 12.10898 | 5.451428 | 85.63081 | 92.70248 | 1485.948 | 594.379 |
| 2.5 | 2 | 1.2 | 5.10519 | 12.76297 | 5.745852 | 90.25562 | 98.74163 | 1650.79 | 660.3161 |
| 2.5 | 2 | 1.4 | 5.382313 | 13.45578 | 6.057753 | 95.15494 | 105.0553 | 1834.873 | 733.9493 |
| 2.5 | 2 | 1.6 | 5.672323 | 14.18081 | 6.384157 | 100.2821 | 111.5968 | 2037.934 | 815.1735 |
| 2.5 | 2 | 1.8 | 5.973159 | 14.9329 | 6.722745 | 105.6006 | 118.3296 | 2259.832 | 903.9329 |
| 2.5 | 2 | 2 | 6.283184 | 15.70796 | 7.071676 | 111.0816 | 125.225 | 2500.505 | 1000.202 |
| 2.5 | 2 | 2.2 | 6.601083 | 16.50271 | 7.429469 | 116.7018 | 132.2595 | 2759.933 | 1103.973 |
| 2.5 | 2 | 2.4 | 6.925785 | 17.31446 | 7.794918 | 122.4423 | 139.4143 | 3038.128 | 1215.251 |
| 2.5 | 2 | 2.6 | 7.256405 | 18.14101 | 8.167029 | 128.2874 | 146.6737 | 3335.118 | 1334.047 |
| 2.5 | 2 | 2.8 | 7.592211 | 18.98053 | 8.544976 | 134.2241 | 154.0248 | 3650.94 | 1460.376 |
| 3 | 2 | 1 | 4.843594 | 14.53078 | 5.451428 | 102.757 | 109.8286 | 2139.764 | 713.2548 |
| 3 | 2 | 1.2 | 5.10519 | 15.31557 | 5.745852 | 108.3067 | 116.7928 | 2377.138 | 792.3793 |
| 3 | 2 | 1.4 | 5.382313 | 16.14694 | 6.057753 | 114.1859 | 124.0863 | 2642.217 | 880.7391 |
| 3 | 2 | 1.6 | 5.672323 | 17.01697 | 6.384157 | 120.3385 | 131.6532 | 2934.625 | 978.2082 |
| 3 | 2 | 1.8 | 5.973159 | 17.91948 | 6.722745 | 126.7207 | 139.4497 | 3254.159 | 1084.72 |
| 3 | 2 | 2 | 6.283184 | 18.84955 | 7.071676 | 133.2979 | 147.4413 | 3600.727 | 1200.242 |
| 3 | 2 | 2.2 | 6.601083 | 19.80325 | 7.429469 | 140.0422 | 155.5999 | 3974.304 | 1324.768 |
| 3 | 2 | 2.4 | 6.925785 | 20.77735 | 7.794918 | 146.9307 | 163.9027 | 4374.905 | 1458.302 |
| 3 | 2 | 2.6 | 7.256405 | 21.76922 | 8.167029 | 153.9448 | 172.3312 | 4802.57 | 1600.857 |
| 3 | 2 | 2.8 | 7.592211 | 22.77663 | 8.544976 | 161.069 | 180.8697 | 5257.353 | 1752.451 |
| 3.5 | 2 | 1 | 4.843594 | 16.95258 | 5.451428 | 119.8831 | 126.9548 | 2912.457 | 832.1306 |
| 3.5 | 2 | 1.2 | 5.10519 | 17.86816 | 5.745852 | 126.3579 | 134.8439 | 3235.549 | 924.4425 |
| 3.5 | 2 | 1.4 | 5.382313 | 18.8381 | 6.057753 | 133.2169 | 143.1173 | 3596.351 | 1027.529 |
| 3.5 | 2 | 1.6 | 5.672323 | 19.85313 | 6.384157 | 140.3949 | 151.7096 | 3994.35 | 1141.243 |
| 3.5 | 2 | 1.8 | 5.973159 | 20.90606 | 6.722745 | 147.8408 | 160.5699 | 4429.271 | 1265.506 |
| 3.5 | 2 | 2 | 6.283184 | 21.99114 | 7.071676 | 155.5142 | 169.6576 | 4900.989 | 1400.283 |
| 3.5 | 2 | 2.2 | 6.601083 | 23.10379 | 7.429469 | 163.3825 | 178.9402 | 5409.469 | 1545.562 |
| 3.5 | 2 | 2.4 | 6.925785 | 24.24025 | 7.794918 | 171.4192 | 188.3912 | 5954.731 | 1701.352 |
| 3.5 | 2 | 2.6 | 7.256405 | 25.39742 | 8.167029 | 179.6023 | 197.9887 | 6536.831 | 1867.666 |
| 3.5 | 2 | 2.8 | 7.592211 | 26.57274 | 8.544976 | 187.9138 | 207.7145 | 7155.842 | 2044.526 |
| 4 | 2 | 1 | 4.843594 | 19.37437 | 5.451428 | 137.0093 | 144.081 | 3804.026 | 951.0064 |
| 4 | 2 | 1.2 | 5.10519 | 20.42076 | 5.745852 | 144.409 | 152.895 | 4226.023 | 1056.506 |
| 4 | 2 | 1.4 | 5.382313 | 21.52925 | 6.057753 | 152.2479 | 162.1482 | 4697.275 | 1174.319 |
| 4 | 2 | 1.6 | 5.672323 | 22.68929 | 6.384157 | 160.4513 | 171.766 | 5217.111 | 1304.278 |
| 4 | 2 | 1.8 | 5.973159 | 23.89263 | 6.722745 | 168.961 | 181.69 | 5785.171 | 1446.293 |
| 4 | 2 | 2 | 6.283184 | 25.13274 | 7.071676 | 177.7306 | 191.8739 | 6401.292 | 1600.323 |
| 4 | 2 | 2.2 | 6.601083 | 26.40433 | 7.429469 | 186.7229 | 202.2806 | 7065.429 | 1766.357 |
| 4 | 2 | 2.4 | 6.925785 | 27.70314 | 7.794918 | 195.9076 | 212.8796 | 7777.608 | 1944.402 |
| 4 | 2 | 2.6 | 7.256405 | 29.02562 | 8.167029 | 205.2598 | 223.6461 | 8537.902 | 2134.475 |
| 4 | 2 | 2.8 | 7.592211 | 30.36884 | 8.544976 | 214.7586 | 234.5593 | 9346.405 | 2336.601 |

2. **The surface distance scale factors,** $\boldsymbol{h}_{\boldsymbol{\theta}}$ **and** $\boldsymbol{h}_{\boldsymbol{\phi}}$ **of Bicelles**

The surface distance scale factors, $h_{\theta}$ and $h_{\phi}$, can be determined directly from the geometry shown in Fig. 1. The distance increment $dS_{\theta}$ along the $\theta$-direction is evaluated in the z-x plane at $\phi$= 0.

${ds}_{\theta}^{2}$=$dz^{2}+dx^{2}$=$\left( \frac{dr}{d\theta}cos\theta d\theta-rcos\theta d\theta\right)^{2}+\left( \frac{dr}{d\theta}sin\theta d\theta+rcos\theta d\theta\right)^{2}$

=$\left( \left( \frac{dr}{d\theta} \right)^{2}+r^{2} \right){d\theta}^{2}$=$\left\{ h_{\theta}\left( \theta\right) \right\}^{2}{d\theta}^{2}$. (S1)

Then,

$h_{\theta}\left( \theta\right)=r\sqrt{\left[ 1+\frac{r^{4}}{4}{sin}^{2}2\theta\left( \frac{1}{b^{2}}-\frac{1}{d^{2}} \right)^{2} \right]}$. (S2)

with

$r= \frac{bd}{\sqrt{\left( b^{2}{sin}^{2}\theta+d^{2}{cos}^{2}\theta\right)}}$ . (S3)

The scale factor $h_{\phi}\left( \phi\right)$ is obtained in the x-y plane by evaluating the distance increment $ds_{\phi}$ along the $\phi$-direction, while keeping $\theta$ fixed.

$ds_{\phi}^{2}=d_{x}^{2}+d_{y}^{2}$=$\left( a+d+rsin\theta\right)^{2}\left( d\phi\right)^{2}$. (S4)

Therefore,

$h_{\phi}\left( \phi\right) =a+d+rsin\theta=R+rsin\theta$. (S5)

It should be noted that the product $h_{\theta}\left( \theta\right)h_{\phi}\left( \phi\right)$ corresponds to the NMR lineshape factor, obtained separately from the Jacobian surface element dS, as defined in Eq (16) of the main text.
